# Dimensions of early life adversity are differentially associated with patterns of delayed and accelerated brain maturation

**DOI:** 10.1101/2024.01.22.576780

**Authors:** Dani Beck, Lucy Whitmore, Niamh MacSweeney, Alexis Brieant, Valerie Karl, Ann-Marie G. de Lange, Lars T. Westlye, Kathryn L. Mills, Christian K. Tamnes

**Author notes:** Correspondence: Dani Beck.

## Abstract

**Background:** Different types of early-life adversity have been associated with children’s brain structure and function. However, understanding the disparate influence of distinct adversity exposures on the developing brain remains a major challenge.

**Methods:** This study investigates the neural correlates of 10 robust dimensions of early-life adversity identified through exploratory factor analysis in a large community sample of youth from the Adolescent Brain Cognitive Development (ABCD) Study. Brain age models were trained, validated, and tested separately on T1-weighted (T1; N = 9524), diffusion tensor (DTI; N = 8834), and resting-state functional (rs-fMRI; N = 8233) magnetic resonance imaging (MRI) data from two time points (mean age = 10.7 years, SD = 1.2, range = 8.9-13.8 years).

**Results:** Bayesian multilevel modelling supported distinct associations between different types of early-life adversity exposures and younger- and older-looking brains. Dimensions generally related to emotional neglect, such as lack of primary and secondary caregiver support, and lack of caregiver supervision, were associated with lower brain age gaps (BAGs), i.e., younger-looking brains. In contrast, dimensions generally related to caregiver psychopathology, trauma exposure, family aggression, substance use and separation from biological parent, and socio-economic disadvantage and neighbourhood safety were associated with higher BAGs, i.e., older-looking brains.

**Conclusions:** The findings suggest that dimensions of early-life adversity are differentially associated with distinct neurodevelopmental patterns, indicative of dimension-specific delayed and accelerated brain maturation.

## 1. Introduction

Early-life adversity (ELA) such as exposure to abuse, violence, neglect, separation from caregivers, and chronic poverty, among others, can have widespread effects on youth neurodevelopment (1) and increase risk for mental disorders (2,3). Previous studies investigating how ELA influences neural development have adopted varied theoretical frameworks, reflecting the complex and multifaceted nature of the field (4,5). Traditionally, research has focused on a single type of adversity; for example, reporting associations between heightened amygdala activation and early exposure to violence (6), and lower volumes of gray matter and low income (7) and exposure to institutionalisation (8). A critical limitation of this approach however, is the fact that most children are exposed to numerous types of adversities concurrently (9). The cumulative risk approach overcomes this by aggregating different forms of adversity into a singular risk factor. Research has found that early cumulative risk was prospectively associated with lower total gray matter volume, cortex volume, and right superior parietal and inferior parietal cortical thickness (10). While the cumulative risk approach is useful for identifying children at greatest risk for intervention and can thus serve as a vital public health tool, aggregating risk factors may obscure potentially diverging effects of different adverse experiences on the developing brain.

More nuanced perspectives that have emerged differentiate between adverse experiences related to threat versus deprivation (11), drawing support for the neural basis of this distinction from studies on fear learning and sensory deprivation. For children exposed to threat, studies have reported lower cortical thickness, surface area, volume of the amygdala and hippocampus (12–14) and ventro-medial prefrontal cortex (vmPFC) (15,16), in addition to reduced resting-state amygdala–vmPFC connectivity (17). The observed effects may be indicative of accelerated maturation and are consistent with life history and evolutionary-biology theories proposing accelerated development as an adaptation to harsh or stressful environments (4,18–20). Similarly, another conceptual model, the stress acceleration hypothesis, argues that adversity may expedite neural development as a means of compensating for the absence of species-expectant maternal buffering of emotional reactivity (21).

For children exposed to deprivation, such as institutional rearing, lack of social and cognitive stimulation, and other forms of parental absence, research has revealed altered structure and function in the frontoparietal network, amygdala-hippocampal-PFC connectivity (11,22), and reductions in cortical gray matter and total brain volume, in addition to widespread cortical thinning (8,23,24). Moreover, electroencephalogram (EEG) studies have shown associations between spectral profiles indicative of delayed patterns of functional and cortical maturation and neglect (25), poverty (26), and parental stress (27).

While previous research has contributed to an understanding of how different features of ELA are associated with unique brain outcomes (28), real-world occurrences of adversity are multifaceted and often co-occur in a complex manner, making it a considerable challenge to precisely account for the heterogeneity in ELA. To address this, several data-driven methods have been applied (29–33), albeit on small or homogenous samples. Recent research by Brieant et al. (2023) capitalised on big data from the Adolescent Brain Cognitive Development (ABCD) Study and identified 10 dimensions of adversity co-occurrence pertaining to conceptual domains reflecting 1) caregiver psychopathology, 2) socioeconomic disadvantage and lack of neighbourhood safety, 3) secondary caregiver lack of support, 4) primary caregiver lack of support, 5) child report of family conflict, 6) caregiver substance use and biological parent separation, 7) family anger and arguments, 8) family aggression, 9) trauma exposure, and 10) caregiver lack of supervision. Yet, to date, the neural correlates of these dimensions of ELA have not been investigated.

Brain age prediction offers a framework that can combine multiple imaging features, alleviating the need for selection of specific metrics or regions, and yields an individualised surrogate marker of brain maturation (35). Brain age involves estimation of biological age based on brain MRI characteristics, which may differ from an individual’s chronological age (36). This difference, termed the brain age gap (BAG), could reflect deviation from typical neurodevelopmental patterns, and has been validated in several neurodevelopmental studies (37–40). Previous literature has also validated the stability of brain age models across early adolescence, with evidence of BAG scores tracking with metrics of maturation (40). Studies have also linked lower BAG in youth to attention-deficit hyperactivity disorder (ADHD), lower socio-economic status, higher anxiety and depression, as well as greater general psychopathology symptom severity (37,41–44). In the context of ELA, where different dimensions of adversity may be associated with unique brain outcomes (4,28,45–49), brain age can probe individualised markers of delayed or accelerated maturation by means of younger- or older-looking brains.

To this end, using ABCD data, our primary aim was to test for associations between MRI-based estimates of brain maturation and 10 previously characterised dimensions of ELA co-occurrence (34). Based on theoretical accounts and the empirical studies reviewed above, we hypothesised ELA dimensions of 1) caregiver psychopathology, 2) socioeconomic disadvantage and lack of neighbourhood safety, 5) child report of family conflict, 6) caregiver substance use and biological parent separation, 7) family anger and arguments, 8) family aggression, and 9) trauma to be associated with older-looking brains and accelerated maturation between the two time points. We hypothesised ELA exposures of 3) secondary caregiver lack of support, 4) primary caregiver lack of support, and 10) caregiver lack of supervision to be associated with younger-looking brains and delayed maturation over time.

## 2. Methods and Materials

### 2.1. Sample and ethical approval

The Adolescent Brain Cognitive Development (ABCD) Study ^®^ (50) comprises of children and adolescents part of an ongoing longitudinal study. Participants were excluded using the ABCD Study exclusion criteria listed elsewhere (51). Data used in the present study were drawn from the ABCD curated annual release 5.0, containing data from baseline up until the second-year visit (https://data-archive.nimh.nih.gov/abcd). All ABCD Study data is stored in the NIMH Data Archive Collection #2573, which is available for registered and authorised users (Request #7474, PI: Westlye). The 5.0 release will be permanently available as a persistent dataset defined in the NDA Study 1299 and has been assigned the DOI 10.15154/8873-zj65. The Institutional Review Board (IRB) at the University of California, San Diego, approved all aspects of the ABCD Study (52). Parents or guardians provided written consent, while the child provided written assent. The current study was conducted in line with the Declaration of Helsinki and was approved by the Norwegian Regional Committee for Medical and Health Research Ethics (REK 2019/943).

### 2.2. Demographic information and data quality assurance

The initial sample consisted of ∼11,800 participants (52% male) at mean age 10.75 (SD = 1.18, range 8.92-13.83) years, with baseline and two-year follow-up observations (obs) of T1 (obs = 19,048), DTI (obs = 17,672), and rs-fMRI (obs = 16,495) data. Quality control procedures followed a standard protocol described in Hagler et al. (2019). Briefly, participants with excessive head motion or poor data quality were excluded from the curated data release by the ABCD Study team. Additional quality assurance was carried out following extraction of data using the recommendations for data cleaning provided by the ABCD Study team (using data structure abcd_imgincl01). Following quality assurance, the final sample included T1 obs of 19,047, DTI obs of 17,668, and rs-fMRI obs of 16,466 used for the current study. Demographic information for each brain MRI modality-specific sample can be found in SI Table 1, while the T1 sample is illustrated in Figure 1.

**Figure 1.**
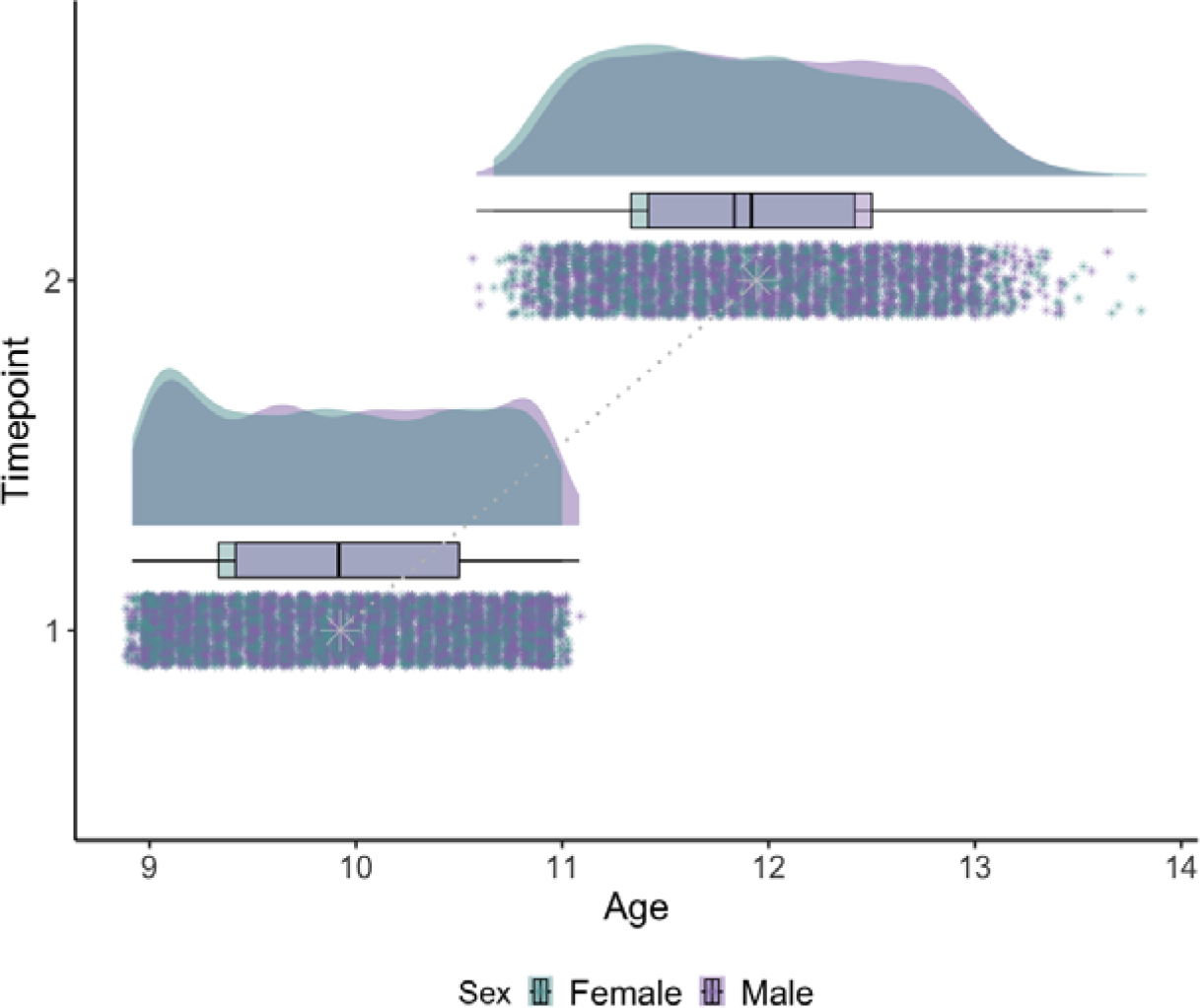
Age distribution split by sex and time point. Time point (1 and 2) represent baseline and two-year-follow-up data from the ABCD Study cohort. Female data in teal blue; male in lavender. Dotted gray line is drawn between mean age at time point 1 and 2.

### 2.3. MRI acquisition, processing, and segmentation

Neuroimaging data were acquired at 21 different sites (using 31 scanners) and processed by the ABCD Study team. A 3-T Siemens Prisma, General Electric 750 or Phillips scanner was used for data acquisition. Protocols used for data acquisition and processing are described in detail elsewhere (50,53) and available in SI Section 1. Three modalities of brain structural and functional measures were used in the present study: structural grey matter measures (T1), diffusion white matter microstructural measures (DTI), and resting-state functional connectivity measures (rs-fMRI) (53). Cortical surface reconstruction and subcortical segmentation was performed with FreeSurfer v7.1.1 (54,55). White matter microstructural measures were generated using AtlasTrack, a probabilistic atlas-based method for automated segmentation of white matter fibre tracts (Hagler et al., 2009). Measures of functional connectivity were computed using a seed-based, correlational approach (57), where average time courses were calculated for cortical surface-based ROIs using a functionally-defined parcellation based on resting-state functional connectivity patterns (58) and subcortical ROIs (55). A detailed description of the processing and extraction of brain imaging data used is provided in full in SI Section 1.

Briefly, For T1, we extracted tabulated total and regional measures of cortical surface area, thickness, volume, sulcal depth, intensity-gray-white contrast, and subcortical volume (397 measures). For DTI, full and inner (multi) shell tissue properties including functional anisotropy (FA) and mean (MD), longitudinal (or axial, AD), and transverse (or radial, RD) diffusivity were extracted for total and regional features (576 measures). For rs-fMRI, functional connectivity within and between parcellations from Gordon network, including subcortical data (58) were extracted (416 measures). Following procedures of quality assurance (Section 2.2), harmonisation of multi-scanner effects was carried out using *longCombat* (59) (see SI Section 2 and SI Figures 1-3).

### 2.4. Brain age prediction

Brain age prediction was carried out using the eXtreme Gradient Boosting (XGBoost) regression model (60), a machine learning algorithm that a gradient boosting library by combining multiple decision trees to create predictive models. Parameters were tuned using ten-fold cross-validation, stratified by age. The models were fitted using the best estimators and optimised models were applied to the (hold-out) test sample. R2, RMSE, and MAE were calculated to evaluate prediction accuracy in the test set. For each brain modality (T1, DTI, rs-fMRI), 50% of the data was used as the hold-out test sample and 50% was used for model training and validation. Here, data was split ensuring an equal distribution of cross-sectional and longitudinal data across training and testing samples, whereby no two datapoints from the same individual (longitudinal) were separated.

Consistent with a recent brain age paper using the ABCD Study sample (61), confounding effects from complex family-related factors were minimised using a group shuffle split with family ID as the group indicator to ensure that no siblings were split across training and test sets. A detailed overview of T1, DTI, and rs-fMRI training and test samples, including demographic information are provided in SI Section 3, SI Table 1, and SI Figures 4-6. A complete list of all the extracted measures used for brain age prediction per modality is provided in SI Tables 2-4. Feature importance scores for each model is provided in SI Figures 7-9. To adjust for commonly observed age-bias (overestimated predictions for younger participants and underestimated predictions for older participants) (62), we applied a statistical correction as previously described in (63). The difference between an individual’s predicted brain age and their chronological age (BAG) was calculated by subtracting the participants chronological age from the age bias corrected predicted age (BAG = predicted age - chronological age) for each of the models, providing T1, DTI, and rs-fMRI-based BAG values for all participants. The resulting BAG can be either positive or negative, indicating an older- or younger-looking brain than the individual’s actual age.

### 2.5. Early life adversity

The current study utilised 10 previously obtained factor scores from 60 measures of early-life adversity (ELA) as detailed in (34). Briefly, Brieant and colleagues identified 139 potential ELA items from the ABCD Study baseline measures, which encompass a spectrum of ELA constructs including caregiving disruption, caregiver psychopathology, maltreatment, neighbourhood safety/violence, among others. These variables were sourced from child and parent reports, as well as researcher assessments, and originated from modified versions of validated scales. To identify dimensions, 60 ELA variables which were binary, polytomous, and continuous in nature were entered into an exploratory factor analysis (EFA) conducted in Mplus version 8.7 (64), resulting in 10 dimensions (F1 to F10; see SI Table 5 and SI Figure 10 for correlation matrix). To obtain factor scores, an exploratory structural equation model was carried out specifying the number of factors identified in the EFA. Further details of the variable selection process, identification of ELA dimensions, and calculation of factor loadings can be found in Brieant et al. (2023).

### 2.6. Statistical analysis

All analyses were carried out using R version 4.2.1 (65). To investigate the association (main effect) between each ELA dimension (F1:F10) and deviation from expected age patterns (i.e., BAG), and whether the effect of each ELA dimension on BAG varies across time points (interaction effect), Bayesian multilevel models were carried out using the *brms* (66,67) R-package. Here, multivariate models are fitted in familiar syntax to comparable frequentist approaches such as a linear mixed effects model using the *lme4* (68). We assessed the relationship between each ELA dimension at baseline and residualised (age-bias corrected) BAG, where modality-specific BAG (T1, DTI, rs-fMRI) was first entered as the dependent variable, each ELA dimension (F1:F10) and interaction term (TP:F1-F10) between time point (TP) and ELA dimensions (F1:F10) were separately entered as independent fixed effects variables with sex entered as a covariate, and with subject ID as the random effect.

To prevent false positives and to regularise the estimated associations, we defined a standard prior around zero with a standard deviation of 0.1 for all regression coefficients, reflecting a baseline expectation of effects being small but allowing for sufficient flexibility in estimation. Each model was run with 8000 iterations, including 4000 warmup iterations, across four chains. This setup was chosen to ensure robust convergence and adequate sampling from the posterior distributions. For each coefficient of interest, we report the mean estimated value of the posterior distribution (b) and its 95% credible interval (the range of values that with 95% confidence contains the true value of the association), and calculated the Bayes Factor (BF) – provided as evidence ratios in the presented figures – using the Savage-Dickey method (69). Briefly, BF can be interpreted as a measure of the strength of evidence (*extreme, very strong, strong, moderate, anecdotal, none*) in favour of the null or alternative hypothesis. For a pragmatic guide on BF interpretation, see SI Table 6.

## 3. Results

### 3.1. Descriptive statistics

Descriptive statistics can be found in SI Table 1. Table 1 summarises descriptive and model validation statistics pertaining to each brain age prediction model, following age-correction. Figure 2 shows predicted age as a function of chronological age for each age prediction model. SI Figure 11 shows a correlation matrix including each modality-specific predicted brain age and BAG.

**Table 1.**
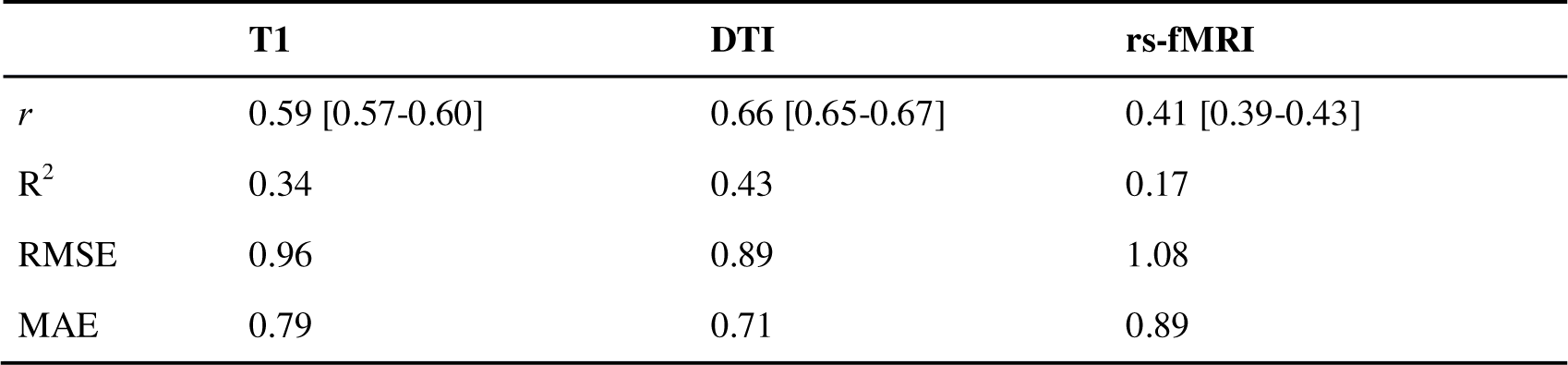
Average R^2^, root mean square error (RMSE), mean absolute error (MAE), and Pearson’s correlations between predicted and chronological age (*r*) for each brain age prediction model, with 95% confidence intervals.

|  | **T1** | **DTI** | **rs-fMRI** |
| --- | --- | --- | --- |
| $r$ | 0.59 [0.57-0.60] | 0.66 [0.65-0.67] | 0.41 [0.39-0.43] |
| $R^2$ | 0.34 | 0.43 | 0.17 |
| RMSE | 0.96 | 0.89 | 1.08 |
| MAE | 0.79 | 0.71 | 0.89 |

**Figure 2.**
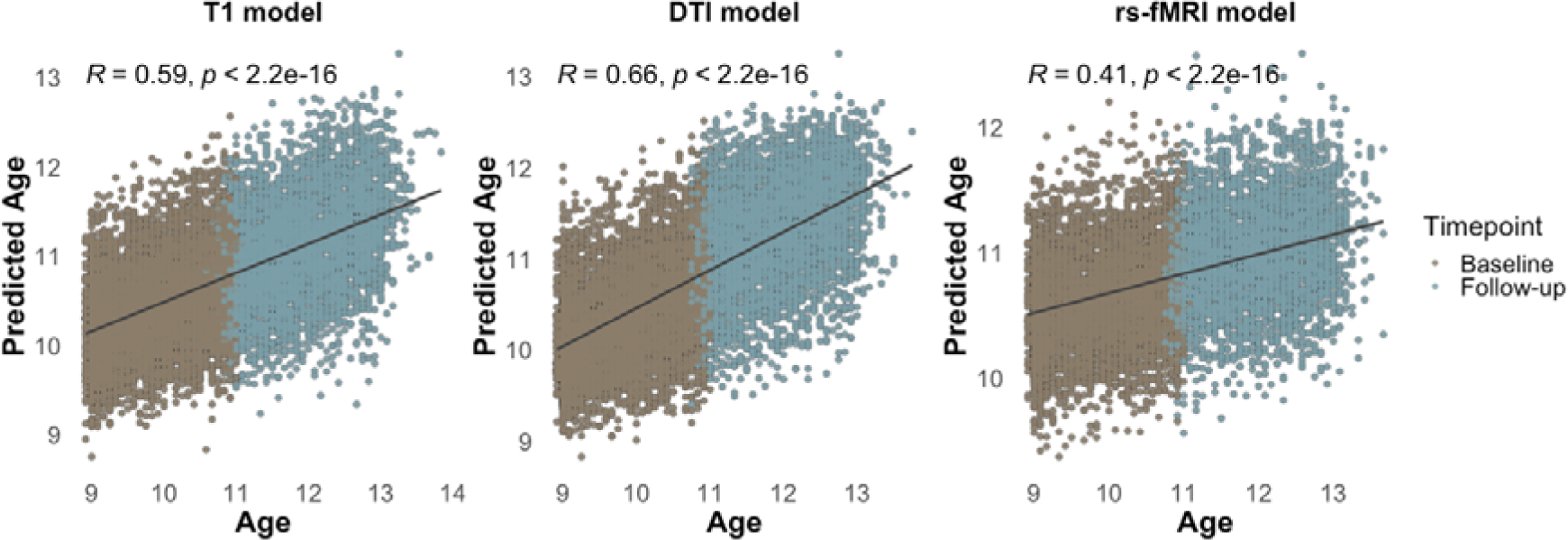
Predicted age as a function of age. Scatter plots demonstrating the correlation between chronological and predicted ages for three brain imaging modalities: T1, DTI, and rs-fMRI. Each plot illustrates the Pearson correlation coefficient (r) and statistical significance (*p*-value), with data points representing individual samples at baseline and follow-up time points. The plots reveal varying degrees of performance accuracy across modalities, with T1 and DTI models showing higher correlations compared to rs-fMRI.

### 3.2. Bayesian multilevel modelling

Bayesian multilevel modelling tested the association between each ELA dimension and modality-specific BAG. The full results are available in SI Tables 7-9 and are visualised below in Figure 3. For estimated credible intervals, see SI Figures 12 and 13.

**Figure 3.**
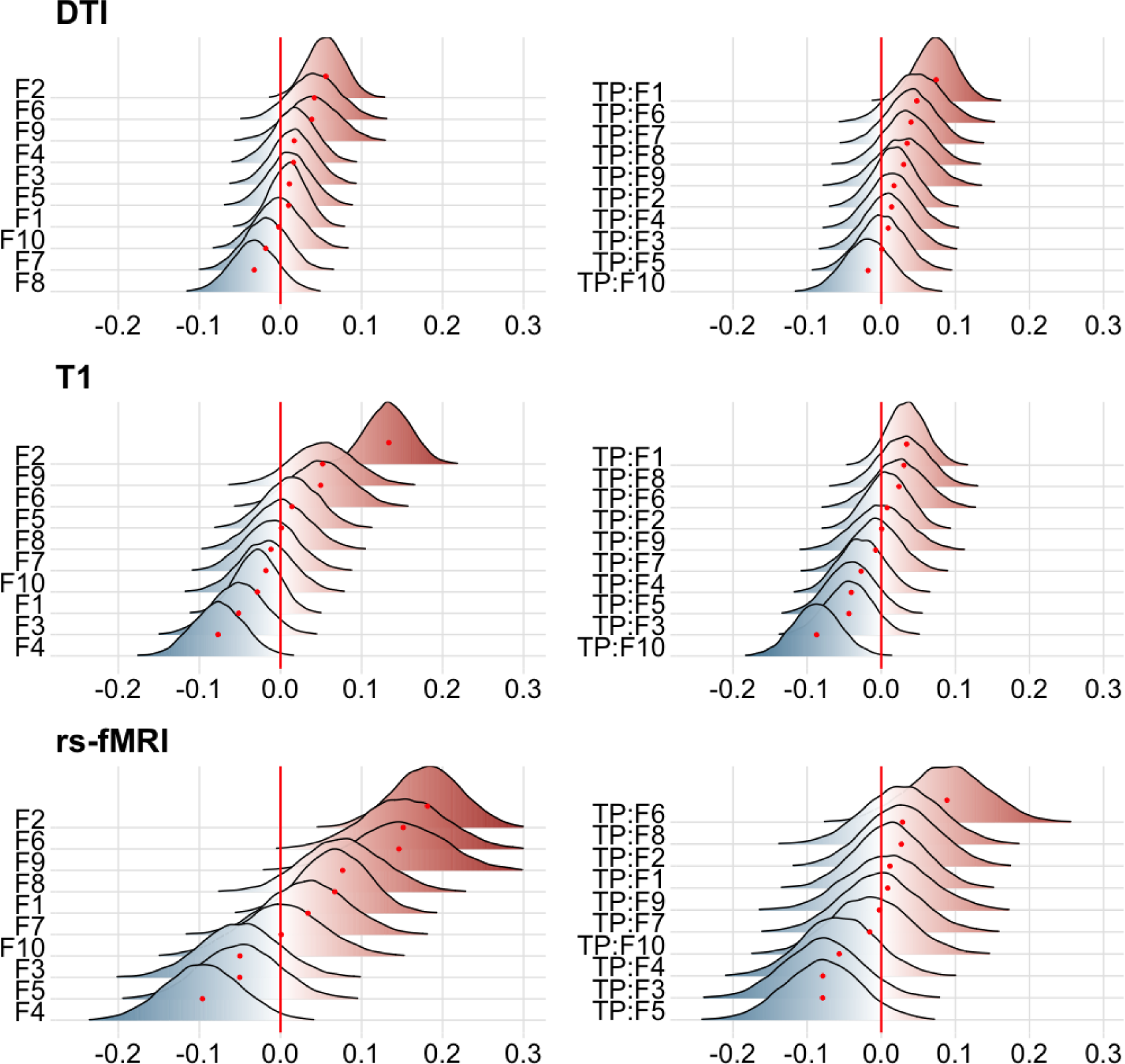
Associations between ELA dimensions and BAG for DTI, T1, and rs-fMRI. The figure shows posterior distributions of the estimates of the standardised coefficient. Estimates for each ELA dimension on BAG (main effect) on the left, and ELA dimension interaction effect of time point (TP:ELA) on BAG on the right. Colour scale follows directionality of evidence, with positive (red) values indicating evidence in favour of positive associations (greater adversity linked to older-looking brains) and negative (blue) values indicating evidence in favour of negative associations (greater adversity linked to younger-looking brains) for each ELA dimension. The width of the distributions represent the uncertainty of the parameter estimates. F1: Caregiver psychopathology; F2: Socio-economic disadvantage and neighbourhood safety; F3: Secondary caregiver lack of support; F4: Primary caregiver lack of support; F5: Youth report of family conflict; F6: Caregiver substance use and separation from biological parent; F7: Family anger and arguments; F8: Family aggression; F9: Trauma exposure; F10: Lack of supervision.

#### 3.2.1. T1 BAG relations

For T1 BAG, the test revealed evidence of a positive association between ELA dimension F2 (socioeconomic disadvantage and neighbourhood safety) and T1 BAG (BF < 0.01, β = 0.13), indicating that this dimension was associated with older-looking brains. Further, the tests revealed evidence of negative associations between ELA dimensions F3 (secondary caregiver lack of support) (BF = 0.9, β = -0.05) and F4 (primary caregiver lack of support) (BF = 0.17, β = -0.08) and T1 BAG, indicating that dimensions related to caregiver emotional neglect are associated with younger-looking brains. In terms of interaction effects of time point, we found a negative association between F10 (lack of supervision) and T1 BAG (BF = 0.08, β = -0.09), indicating that unsupervised youth diverge more from normative age patterns throughout the course of the study period.

#### 3.2.2. DTI BAG relations

For DTI, the tests revealed evidence of a positive association between F2 and DTI BAG (BF = 0.27, β = 0.06), aligning with the T1 BAG findings, and indicating that those living in more disadvantaged and less safe environments may have older-looking brains. In terms of interaction effects of time point, the tests revealed evidence supporting a positive association between ELA dimension F1 (caregiver psychopathology) and DTI BAG (BF = 0.09, β = 0.07), indicating that the brain ages of these youth will diverge more (accelerate) from normative age patterns over time.

#### 3.2.3. rs-fMRI BAG relations

For rs-fMRI, the tests revealed evidence of a positive association between F2 and rs-fMRI BAG (BF < 0.01, β = 0.18), aligning with findings from DTI and T1 BAG. The tests also revealed positive associations between F1 (BF = 0.63, β = 0.07), F6 (caregiver substance abuse and separation from biological parents) (BF = 0.03, β = 0.15), F8 (family aggression) (BF = 0.66, β = 0.08), and F9 (trauma exposure) (BF = 0.07, β = 0.15) and rs-fMRI BAG. These positive associations largely indicate that dimensions linked to a threatening environment relate to older-looking brains. Further, the tests revealed a negative association between F4 and rs-fMRI BAG (BF = 0.27, β = -0.10), in line with findings from T1 BAG. In terms of interaction effects of time point, we found evidence supporting negative associations between both F3 (BF = 0.62, β = -0.08) and F5 (family conflict) (BF = 0.58, β = -0.08) and rs-fMRI BAG, and a positive association between F6 (BF = 0.51, β = 0.09) and rs-fMRI BAG.

## 4. Discussion

Research indicates that approximately half of all children will experience at least one form of adversity by the time they reach adulthood (2,70). Different forms of early-life adversity co-occur and may uniquely impact child brain structure and function. This work sought to delineate the potentially unique brain outcomes of 10 co-occurring dimensions of early-life adversity in a longitudinal sample of 9–14-year-old youth. Our main findings indicate that ELA dimensions of caregiver lack of support and supervision are associated with younger-looking brains and that ELA dimensions of caregiver psychopathology, socioeconomic disadvantage and neighbourhood safety, caregiver substance abuse and separation from biological parents, family aggression, and trauma exposure are associated with older-looking brains. Our findings largely align with and extend the current literature on the differential impact of different co-occurring patterns of environmental exposures and adversity on brain development (45,49).

### 4.1. Links to accelerated brain maturation

Families living in lower socioeconomic status neighbourhoods are exposed to more harms, such as interpersonal violence (71), and are more likely to have concerns about neighbourhood safety (72). Positive associations between the dimension representing socioeconomic disadvantage and neighbourhood safety (F2) and BAG were found with all three brain age models, and are consistent with research showing socioeconomic disadvantage and neighbourhood violence linked to smaller cortical volumes and greater cortical thinning (73–75). Positive associations between exposure to trauma (F9) and BAG is also consistent with previous research showing that children exposed to trauma are more likely to be misclassified as adults by means of older DNA methylation age compared to chronological age, and earlier pubertal maturation (76–78).

For caregiver psychopathology (F1), we found both positive BAG associations as main effects and as interaction effects of time, suggesting that parent psychopathology-related deviations from expected age-patterns accelerate over time. Our findings are challenging to interpret in the context of mixed results from previous research. ABCD Study findings have revealed smaller volume in the right putamen (79), left hippocampus (80), and bilateral hippocampi (81) in relation to parental psychopathology. These findings reflect age patterns of hippocampi and putamen usually not seen until late adolescents and young adulthood – whereby subcortical volumes are expected to reduce over time (82,83) – suggesting the results may indicate accelerated ageing in this younger sample, and thus aligning with the directionality of our results. However, studies using other datasets have found that parental history of depression is associated with larger volume in the bilateral amygdala (84), and others have reported no effects (85). ELA dimensions that indicated the potential presence of household hostility including family aggression (e.g., throwing things, hitting) (F8) revealed a positive main effect for rs-fMRI BAG, but no effects for family arguments (e.g., expressing anger, fighting, raising voices) (F7), with the latter finding going against our hypothesis.

Co-occurrence of caregiver substance abuse and separation from biological parents in one dimension (F6) might reflect child custody issues related to caregiver substance use disorders or arrests (34,86). Moreover, this dimension also has factor loadings from domestic violence. Our results revealed positive rs-fMRI BAG associations both as a main effect and an interaction effect of time. Drawing from concepts of stress acceleration, parental deprivation accelerates the functional development of the mPFC in children, such that amygdala–mPFC interactions are more adultlike following deprivation experiences (21). Children in this group may have accelerated maturation as a means of adapting from a state of parent-regulated to self-regulated emotional processing due to absent or inconsistent parental care (4,19–21). Further research is needed to understand the underlying mechanisms at play. In summary, dimensions of ELA co-occurrence related to older-looking brains support research reporting accelerated brain maturation in children exposed to potentially more hostile or dangerous environments.

### 4.2. Links to delayed brain maturation

Neurobiological studies of brain development have long assumed a deficit model in which lack of input to the development of a child will result in delay of certain skills (87). In the child brain, this may be reflected by a delay in pruning and thus larger brain volumes and younger-looking brains. Our results support this, with lack of primary (F4) and secondary (F3) caregiver support as well as lack of caregiver supervision (F10) all revealing negative associations with T1 and rs-fMRI BAG, with main effects for F3 and F4, and interaction effects for F3 and F10. Importantly, these factors each share elements of emotional neglect.

Previous research investigating more severe forms of neglect (physical and emotional) have found that children reared in institutions demonstrate EEG patterns suggestive of a delay in cortical maturation in frontal, temporal, and occipital regions (25). However, a wealth of studies also report conflicting results indicative of advanced maturation in similar samples (8,24). A caveat of research carried out on neglect is that children are often in environments enriched for several co-occurring ELAs, making it difficult to rely on the stability of the neglect scores while considering other stressors. Importantly, our results specifically capture emotional forms of neglect in terms of lack of household emotional support in a community with relatively lower levels of risk, which may explain divergence from prior studies on extreme physical neglect or institutionalisation.

Lastly, a negative interaction effect of time for rs-fMRI BAG and family conflict (F5) was also found, indicating that youth exposed to family conflict diverge more from normative age patterns (i.e., delayed maturation) over time. This contradicted our hypothesis and findings from previous research also utilising the ABCD Study cohort which reported high family conflict associations with smaller cortical surface areas of the orbitofrontal cortex, anterior cingulate cortex, and middle temporal gyrus (88). In summary, ELA dimensions related to emotional neglect were associated with younger-looking brains and delayed maturation.

### 4.3. Strengths and limitations

Our study adds new insight into the neural correlates of co-occurring ELA dimensions in a large-scale longitudinal community sample. Using a sample not enriched for adversity exposure has the added value of demonstrating that even less severe ELA exposure is associated with changes in the developing brain, facilitating broader generalisation. However, there is trade-off in that our findings cannot necessarily be generalised to interpret neural correlates of children exposed to more severe forms of adversity, as these groups may not be well-represented in the ABCD Study. Future research is required to address this without losing adequate power. There remain also additional challenges such as accounting for differences in chronicity of adversity events, interindividual differences in resilience, and overlap in adversity types.

Building on data driven methods applied in Brieant et al. (2023), the current study benefitted from ELA dimensions accounting for more variability in ELA patterns and a dimensional structure replicated in an independent sample, suggesting some stability. The current study utilised three estimates of brain age based on different MRI modalities in an attempt to capture potentially tissue-specific effects of ELA dimensions. While the multimodal (with longitudinal data) approach is a strength, deep learning methods have shown greater accuracy in recent years (39,89). Moreover, focusing on different regional rather than global brain metrics and their association with adversity dimensions represents an opportunity for future research.

### 4.4. Conclusion

The current study supports notions that brain MRI outcomes related to early-life adversity are differentially associated with accelerated and delayed brain maturation. Neurodevelopmental processes influenced by experiences of trauma, parental psychopathology, socio-economic disadvantage and neighbourhood safety, caregiver substance abuse and separation from biological parents, and family aggression, are at least partially distinct from those influenced by experiences of emotional neglect, with brain age deviations indicating differential maturational patterns. Future research should build on this work by investigating, for example, how brain age patterns mediate the associations between dimensions of early life adversity and child behavioural and symptom measures. Such studies could elucidate whether indices of brain maturation serve as an underlying biological mechanism linking early adversity to later child outcomes, thereby offering a more comprehensive understanding of these developmental processes and helping guide intervention strategies that aim to mitigate the impact of early-life adversity and supporting healthy development of at-risk youth.

## 6. Acknowledgments

Data used in the preparation of this article were obtained from the Adolescent Brain Cognitive DevelopmentSM (ABCD) Study (https://abcdstudy.org), held in the NIMH Data Archive (NDA). This is a multisite, longitudinal study designed to recruit more than 10,000 children aged 9-10 and follow them over 10 years into early adulthood. The ABCD Study® is supported by the National Institutes of Health and additional federal partners under award numbers U01DA041048, U01DA050989, U01DA051016, U01DA041022, U01DA051018, U01DA051037, U01DA050987, U01DA041174, U01DA041106, U01DA041117, U01DA041028, U01DA041134, U01DA050988, U01DA051039, U01DA041156, U01DA041025, U01DA041120, U01DA051038, U01DA041148, U01DA041093, U01DA041089, U24DA041123, U24DA041147. A full list of supporters is available at https://abcdstudy.org/federal-partners.html. A listing of participating sites and a complete listing of the study investigators can be found at https://abcdstudy.org/consortium_members/. ABCD consortium investigators designed and implemented the study and/or provided data but did not necessarily participate in the analysis or writing of this report. This manuscript reflects the views of the authors and may not reflect the opinions or views of the NIH or ABCD consortium investigators. This manuscript has been posted on the preprint server *bioRxiv*: https://www.biorxiv.org/content/10.1101/2024.01.22.576780v1. We thank the co-authors of Brieant et al. (2023) for their contributions to the development of the factor scores used in this paper (Anna Vannucci, Hajer Nakua, Jenny Harris, Jack Lovell, Divya Brundavanam, Nim Tottenham, and Dylan Gee).

