## Supplementary Information for "Dimensions of early life adversity are differentially associated with patterns of delayed and accelerated brain maturation"

**SI Section 1**. MRI acquisition and processing

For T1-weighted MRI acquisitions on Siemens and Philips 3T scanners, a matrix of 256 × 256, 176 slices (Siemens) and 225 slices (Philips) with a field of view (FOV) of 256 × 256, echo time (TE)/repetition time (TR) (ms) of 2.88/2500 (Siemens) and 2.9/6.31 (Philips), flip angle of 8°. On GE scanners, the same matrix and FOV were used with TE/TR (ms) of 2/2500 and a flip angle of 8°. The spatial resolution was consistent at 1.0 × 1.0 × 1.0 mm across all three platforms. The dMRI acquisition (1.7 mm isotropic) uses multiband EPI with slice acceleration factor 3 and includes 96 diffusion directions, seven b = 0 frames, and four b-values (6 directions with b = 500 s/mm^2^, 15 directions with b = 1000 s/mm^2^, 15 directions with b = 2000 s/mm^2^, and 60 directions with b = 3000 s/mm^2^). For rs-fMRI acquisition, the following scanning parameters were used: matrix of 90 × 90, 60 slices, FOV = 216 × 216, TE/TR = 800/30, flip angle = 52° and resolution = 2.4 × 2.4 × 2.4 mm. The fMRI acquisitions (2.4 mm isotropic, TR = 800 ms) used multiband EPI with slice acceleration factor 6 and were split into 2-4 5-minute scanning sessions, in which participants were instructed to keep their eyes open and fixate on a crosshair.

For T1-weighted MRI data, cortical surface reconstruction and subcortical segmentation was performed with FreeSurfer v7.1.1 (Dale et al., 1999; Fischl et al., 2002). This processing includes motion correction and averaging (Reuter et al., 2010), removal of non-brain tissue (Ségonne et al., 2004), automated Talairach transformation, segmentation of the subcortical white matter and deep gray matter volumetric structures (Fischl et al., 2002, 2004), intensity normalization (Sled et al., 1998), tessellation of the gray matter white matter boundary, automated topology correction (Fischl et al., 2001; Ségonne et al., 2007), and surface deformation (Dale et al., 1999). From the ABCD Data Repository, we obtained tabulated data including total and regional measures of cortical surface area, thickness, volume, sulcal depth, intensity-gray-white contrast, and subcortical volume (397 measures) from the following data structures: mri_y_smr_thk_dsk, mri_y_smr_sulc_dsk, mri_y_smr_area_dsk, mri_y_smr_t1_contr_dsk, mri_y_smr_vol_dsk, mri_y_smr_vol_aseg.

For DTI data, processing was carried out using AtlasTrack, a probabilistic atlas-based method for automated segmentation of white matter fibre tracts (Hagler et al., 2009). A detailed pipeline can be found in Hagler et al. (2019). Briefly, diffusion tensor parameters are calculated using a standard, linear estimation approach with log-transformed diffusion-weighted (DW) signals (Basser et al., 1994), whereby two tensor models are fitted. In the first DTI model fit (DTI inner shell, or DTI_IS_), frames with b > 1000 s/mm^2^ are excluded from tensor fitting (leaving 6 directions at b = 500 s/mm^2^ and 15 directions at b = 1000 s/mm^2^) so that the derived diffusivity measures better correspond to those from traditional, single-b-value acquisitions. In the second DTI model fit (DTI full shell, or DTI_FS_), all gradient strengths/shells (6 directions at b = 500 s/mm^2^, 15 directions at b = 1000 s/mm^2^, 15 directions at b = 2000 s/mm^2^, and 60 directions at b = 3000 s/mm^2^) are included. For the current study, both full and inner shell tissue properties of functional anisotropy (FA) and mean (MD), longitudinal (or axial, AD), and transverse (or radial, RD) diffusivity were extracted for total and regional features (576 measures). This involved obtaining the following tabulated data structures from the ABCD Data Repository: mri_y_dti_fa_fs_at, mri_y_dti_fa_fs_aseg, mri_y_dti_ld_fs_at, mri_y_dti_ld_fs_aseg, mri_y_dti_md_fs_at, mri_y_dti_md_fs_aseg, mri_y_dti_td_fs_at, mri_y_dti_td_fs_aseg, mri_y_dti_fa_is_at, mri_y_dti_fa_is_aseg, mri_y_dti_ld_is_at, mri_y_dti_ld_is_aseg, mri_y_dti_md_is_at, mri_y_dti_md_is_aseg, mri_y_dti_td_is_at, mri_y_dti_td_is_aseg.

For fMRI, preprocessing steps are outlined in detail in Hagler et al. (2019). Briefly, head motion is corrected by registering each frame to the first using AFNI’s 3dvolreg (Cox, 1996), while B_0_ distortions are corrected using the same reversing polarity method used for the dMRI (Holland et al., 2010). Preprocessing also includes steps to avoid signal drop-out, corrections for distortions due to gradient nonlinearities (Jovicich et al., 2006) and between-scan motion correction. Additional rs-fMRI processing steps included removal of initial frames, normalization, regression, temporal filtering, and calculation of ROI-average time courses. Measures of functional connectivity were computed using a seed-based, correlational approach (Van Dijk et al., 2010), where average time courses were calculated for cortical surface-based ROIs using a functionally-defined parcellation based on resting-state functional connectivity patterns (Gordon et al., 2016) and for subcortical ROIs (Fischl et al., 2002).

Correlation coefficients between the average time courses of each ROI-pairing were then calculated. Here, correlations between unique pairs of ROIs are obtained and Fisher transformed into z-statistics. Connectivity measures represent the averaged Fisher-transformed correlations of all the unique pairs of ROIs either within a cortical network, between cortical networks, or between cortical networks and subcortical regions. ROIs were grouped into Gordon networks and within and between network correlation strength was extracted by calculating the mean correlation coefficient of the respective ROI-pairings. Together with the correlations between the average time course within a network and each subcortical ROI, these coefficients served as functional connectivity measures for the current study. From the ABCD Data Repository, we obtained functional connectivity within and between parcellations from the Gordon network, including subcortical data (416 measures) from the following data structures: mri_y_rsfmr_cor_gp_aseg and mri_y_rsfmr_cor_gp_gp.

Quality assurance procedures carried out on all outlined brain features for each modality are outlined in section 2.2 of the main manuscript.

**SI Section 2**. LongCombat

The ABCD Study data is collected from 21 different sites in the U.S, and children are scanned using thirty-one different magnetic resonance imaging (MRI) scanners from three manufacturing brands (3-T Siemens Prisma, General Electric 750 or Phillips). Due to the technical variability of using multiple scanners, noise and bias can be introduced into the estimation of biological features of interest. Longitudinal ComBat is a powerful harmonisation procedure for longitudinal datasets and controls for type I error better than unharmonized data with scanner included as a covariate (Beer et al., 2020). In the current study, LongCombat was implemented on baseline and two-year follow-up observations (obs) of T1 (obs = 19,047), DTI (obs = 17,668), and rs-fMRI (obs = 16,466) data from the ABCD Study cohort, with each MRI modality being harmonised separately, including covariates of age, sex, and timepoint following recommendations from Beer et al. (2020). SI Figures 1-3 shows MRI data from each modality for selected regions prior to and post harmonisation.

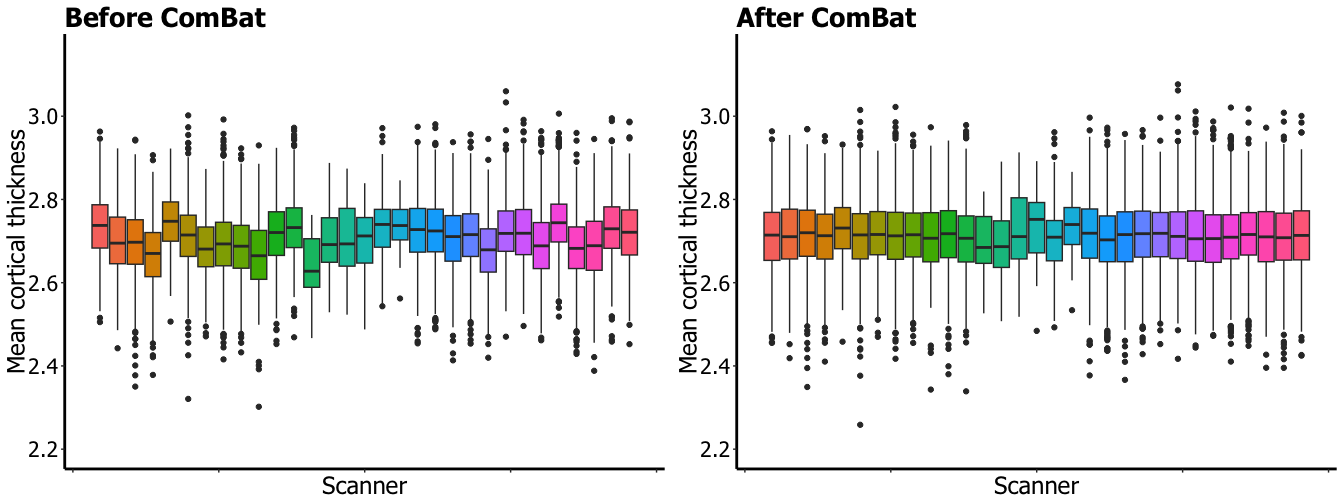

**SI Figure 1.** Distributions of mean cortical thickness in mm for whole brain (smri_thick_cdk_mean) across scanners before and after LongCombat.

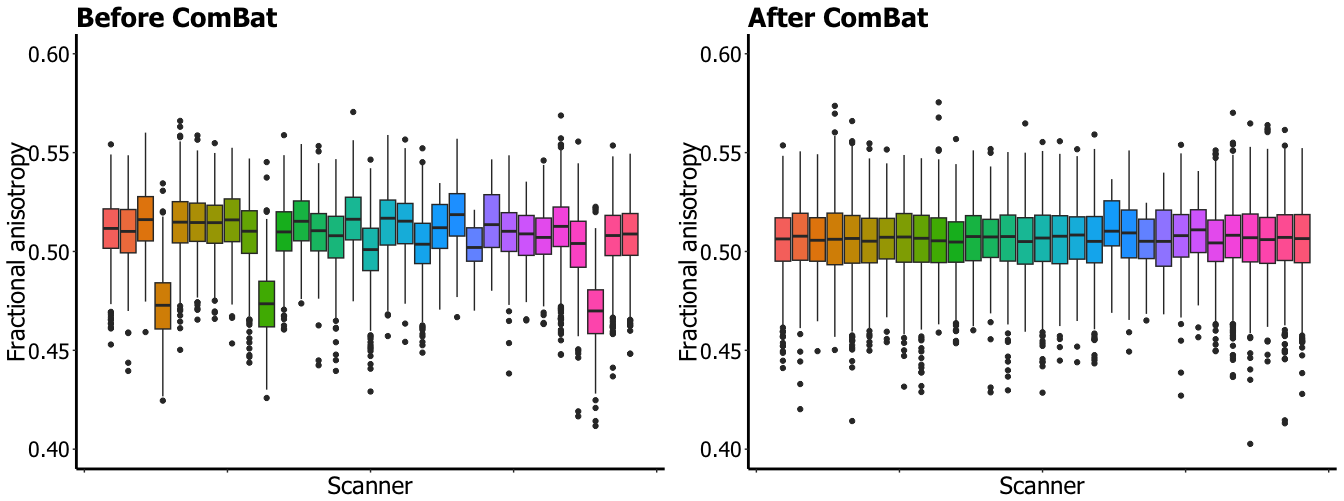

**SI Figure 2.** Distributions of average fractional anisotropy within all DTI atlas tract fibers (dmri_dtifa_fiberat_allfibers.combat) across scanners before and after LongComBat.

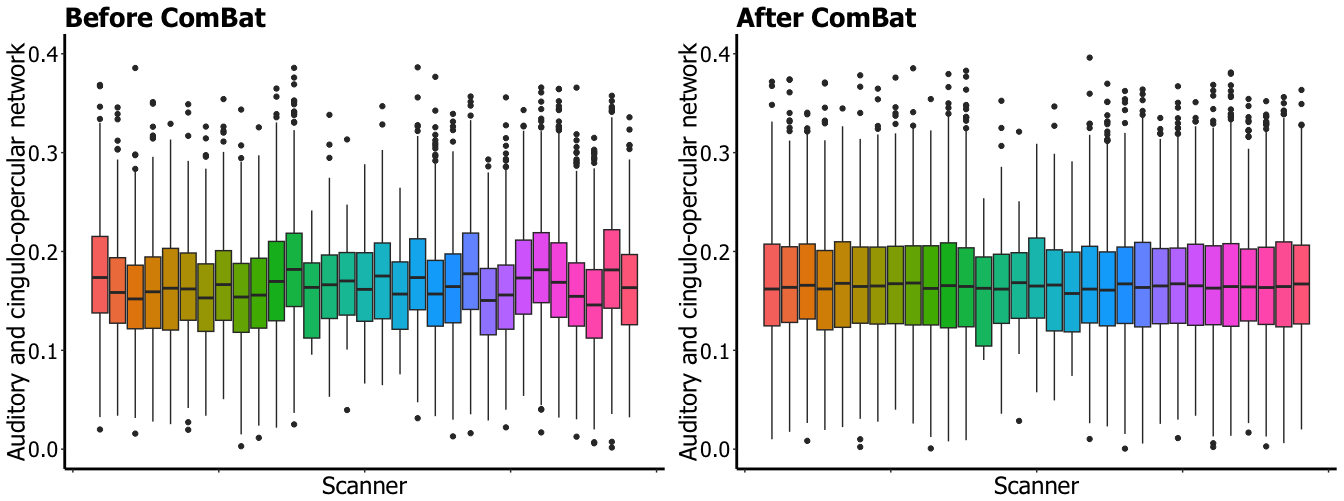

**SI Figure 3.** Distributions of average correlation between auditory network and cingulo-opercular network (rsfmri_c_ngd_ad_ngd_cgc) across scanners before and after LongComBat.

**SI Section 3.** Brain age prediction training and test sample overview

Our imaging data selection was informed by prior brain age studies using T1, dMRI, and fMRI data (Beck, de Lange, Alnæs, et al., 2022; Beck, de Lange, Pedersen, et al., 2022; Beck et al., 2021; de Lange et al., 2020; Lund et al., 2022; Subramaniapillai et al., 2022; Voldsbekk et al., 2021). Although T1 and dMRI models are more common, we included rs-fMRI due to its extensive use in early life adversity research. This allows us to examine tissue-specific effects across imaging types. We chose validated metrics from previous studies: fractional anisotropy, diffusivity measures for DTI; cortical surface area, thickness, subcortical volume for T1; and functional connectivity for rs-fMRI. This yielded 416 features for rs-fMRI and 576 for DTI. To match feature counts and reduce bias, we added intensity-gray-white contrast and sulcal depth to the T1 model, increasing its features to 397. For each brain modality (T1, DTI, rs-fMRI), 50% of the data was used as the hold-out test sample and 50% was used for the model training and validation, split 80% and 20% respectively. The sample sizes of each modality are provided in SI Table 1 below. Three Welch two sample t-tests carried out to test the differences in age between the training and test sample across the different models revealed no statistically significant difference for T1 (t = 0.52, df = 19,045, *p* = 0.61), DTI (t = 0.07, df = 17,665, *p* = 0.94), and rs-fMRI (t = 0.30, df = 16,464, *p* = 0.76). The age distribution for each brain MRI modality is provided below (SI Figures 4-6).

| **SI Table 1.** Training and test sample demographic information. | | | | | | | |
| --- | --- | --- | --- | --- | --- | --- | --- |
| **T1** |  | **Training** *(N = 9524)* | | | **Test** *(N = 9523)* | | |
|  |  | *TP1* | *TP2* | *Total* | *TP1* | *TP2* | *Total* |
|  | *Age mean* | 9.91 | 11.93 | 10.73 | 9.92 | 11.94 | 10.73 |
|  | *Age SD* | 0.62 | 0.64 | 1.17 | 0.63 | 0.65 | 1.18 |
|  | *Age range* | 8.92-11.00 | 10.75-13.75 | 8.91-13.75 | 8.92-11.08 | 10.58-13.83 | 8.92-13.83 |
|  | *Sex (M)* |  |  | 4960 (52.1%) |  |  | 5073 (53.3%) |
|  | *Sex (F)* |  |  | 4564 (47.9%) |  |  | 4450 (46.4%) |
| **DTI** |  | **Training** *(N = 8834)* | | | **Test** *(N = 8834)* | | |
|  |  | *TP1* | *TP2* | *Total* | *TP1* | *TP2* | *Total* |
|  | *Age mean* | 9.92 | 11.94 | 10.76 | 9.93 | 11.94 | 10.76 |
|  | *Age SD* | 0.63 | 0.66 | 1.18 | 0.62 | 0.64 | 1.17 |
|  | *Age range* | 8.92-11.08 | 10.58-13.83 | 8.92-13.83 | 8.92-11.00 | 10.58-13.75 | 8.92-13.75 |
|  | *Sex (M)* |  |  | 4626 (52.4%) |  |  | 4638 (52.5%) |
|  | *Sex (F)* |  |  | 4208 (47.6%) |  |  | 4196 (47.5%) |
| **rsfMRI** |  | **Training** *(N = 8233)* | | | **Test** *(N = 8233)* | | |
|  |  | *TP1* | *TP2* | *Total* | *TP1* | *TP2* | *Total* |
|  | *Age mean* | 9.94 | 11.95 | 10.78 | 9.93 | 11.94 | 10.78 |
|  | *Age SD* | 0.63 | 0.65 | 1.18 | 0.63 | 0.65 | 1.18 |
|  | *Age range* | 8.92-11.00 | 10.58-13.83 | 8.92-13.83 | 8.92-11.08 | 10.67-13.67 | 8.92-13.67 |
|  | *Sex (M)* |  |  | 4286 (52.1%) |  |  | 4187 (50.9%) |
|  | *Sex (F)* |  |  | 3947 (47.9%) |  |  | 4046 (49.1%) |
| ***Note****: TP1 = baseline data, TP2 = follow-up data, SD = Standard deviation, M = Male, F = Female.* | | | | | | | |

**SI Table 2**. Showing list of T1-weighted tabulated brain features (N = 397) extracted from the ABCD Study that was used for brain age prediction.

smri_thick_cdk_banksstslh, smri_thick_cdk_cdacatelh, smri_thick_cdk_cdmdfrlh, smri_thick_cdk_cuneuslh, smri_thick_cdk_ehinallh, smri_thick_cdk_fusiformlh, smri_thick_cdk_ifpllh, smri_thick_cdk_iftmlh, smri_thick_cdk_ihcatelh, smri_thick_cdk_locclh, smri_thick_cdk_lobfrlh, smri_thick_cdk_linguallh, smri_thick_cdk_mobfrlh, smri_thick_cdk_mdtmlh, smri_thick_cdk_parahpallh, smri_thick_cdk_paracnlh, smri_thick_cdk_parsopclh, smri_thick_cdk_parsobislh, smri_thick_cdk_parstgrislh, smri_thick_cdk_pericclh, smri_thick_cdk_postcnlh, smri_thick_cdk_ptcatelh, smri_thick_cdk_precnlh, smri_thick_cdk_pclh, smri_thick_cdk_rracatelh, smri_thick_cdk_rrmdfrlh, smri_thick_cdk_sufrlh, smri_thick_cdk_supllh, smri_thick_cdk_sutmlh, smri_thick_cdk_smlh, smri_thick_cdk_frpolelh, smri_thick_cdk_tmpolelh, smri_thick_cdk_trvtmlh, smri_thick_cdk_insulalh, smri_thick_cdk_banksstsrh, smri_thick_cdk_cdacaterh, smri_thick_cdk_cdmdfrrh, smri_thick_cdk_cuneusrh, smri_thick_cdk_ehinalrh, smri_thick_cdk_fusiformrh, smri_thick_cdk_ifplrh, smri_thick_cdk_iftmrh, smri_thick_cdk_ihcaterh, smri_thick_cdk_loccrh, smri_thick_cdk_lobfrrh, smri_thick_cdk_lingualrh, smri_thick_cdk_mobfrrh, smri_thick_cdk_mdtmrh, smri_thick_cdk_parahpalrh, smri_thick_cdk_paracnrh, smri_thick_cdk_parsopcrh, smri_thick_cdk_parsobisrh, smri_thick_cdk_parstgrisrh, smri_thick_cdk_periccrh, smri_thick_cdk_postcnrh, smri_thick_cdk_ptcaterh, smri_thick_cdk_precnrh, smri_thick_cdk_pcrh, smri_thick_cdk_rracaterh, smri_thick_cdk_rrmdfrrh, smri_thick_cdk_sufrrh, smri_thick_cdk_suplrh, smri_thick_cdk_sutmrh, smri_thick_cdk_smrh, smri_thick_cdk_frpolerh, smri_thick_cdk_tmpolerh, smri_thick_cdk_trvtmrh, smri_thick_cdk_insularh, smri_thick_cdk_meanlh, smri_thick_cdk_meanrh, smri_thick_cdk_mean, smri_sulc_cdk_banksstslh, smri_sulc_cdk_cdacatelh, smri_sulc_cdk_cdmdfrlh, smri_sulc_cdk_cuneuslh, smri_sulc_cdk_ehinallh, smri_sulc_cdk_fusiformlh, smri_sulc_cdk_ifpllh, smri_sulc_cdk_iftmlh, smri_sulc_cdk_ihcatelh, smri_sulc_cdk_locclh, smri_sulc_cdk_lobfrlh, smri_sulc_cdk_linguallh, smri_sulc_cdk_mobfrlh, smri_sulc_cdk_mdtmlh, smri_sulc_cdk_parahpallh, smri_sulc_cdk_paracnlh, smri_sulc_cdk_parsopclh, smri_sulc_cdk_parsobislh, smri_sulc_cdk_parstgrislh, smri_sulc_cdk_pericclh, smri_sulc_cdk_postcnlh, smri_sulc_cdk_ptcatelh, smri_sulc_cdk_precnlh, smri_sulc_cdk_pclh, smri_sulc_cdk_rracatelh, smri_sulc_cdk_rrmdfrlh, smri_sulc_cdk_sufrlh, smri_sulc_cdk_supllh, smri_sulc_cdk_sutmlh, smri_sulc_cdk_smlh, smri_sulc_cdk_frpolelh, smri_sulc_cdk_tmpolelh, smri_sulc_cdk_trvtmlh, smri_sulc_cdk_insulalh, smri_sulc_cdk_banksstsrh, smri_sulc_cdk_cdacaterh, smri_sulc_cdk_cdmdfrrh, smri_sulc_cdk_cuneusrh, smri_sulc_cdk_ehinalrh, smri_sulc_cdk_fusiformrh, smri_sulc_cdk_ifplrh, smri_sulc_cdk_iftmrh, smri_sulc_cdk_ihcaterh, smri_sulc_cdk_loccrh, smri_sulc_cdk_lobfrrh, smri_sulc_cdk_lingualrh, smri_sulc_cdk_mobfrrh, smri_sulc_cdk_mdtmrh, smri_sulc_cdk_parahpalrh, smri_sulc_cdk_paracnrh, smri_sulc_cdk_parsopcrh, smri_sulc_cdk_parsobisrh, smri_sulc_cdk_parstgrisrh, smri_sulc_cdk_periccrh, smri_sulc_cdk_postcnrh, smri_sulc_cdk_ptcaterh, smri_sulc_cdk_precnrh, smri_sulc_cdk_pcrh, smri_sulc_cdk_rracaterh, smri_sulc_cdk_rrmdfrrh, smri_sulc_cdk_sufrrh, smri_sulc_cdk_suplrh, smri_sulc_cdk_sutmrh, smri_sulc_cdk_smrh, smri_sulc_cdk_frpolerh, smri_sulc_cdk_tmpolerh, smri_sulc_cdk_trvtmrh, smri_sulc_cdk_insularh, smri_sulc_cdk_meanlh, smri_sulc_cdk_meanrh, smri_sulc_cdk_mean, smri_area_cdk_banksstslh, smri_area_cdk_cdacatelh, smri_area_cdk_cdmdfrlh, smri_area_cdk_cuneuslh, smri_area_cdk_ehinallh, smri_area_cdk_fusiformlh, smri_area_cdk_ifpllh, smri_area_cdk_iftmlh, smri_area_cdk_ihcatelh, smri_area_cdk_locclh, smri_area_cdk_lobfrlh, smri_area_cdk_linguallh, smri_area_cdk_mobfrlh, smri_area_cdk_mdtmlh, smri_area_cdk_parahpallh, smri_area_cdk_paracnlh, smri_area_cdk_parsopclh, smri_area_cdk_parsobislh, smri_area_cdk_parstgrislh, smri_area_cdk_pericclh, smri_area_cdk_postcnlh, smri_area_cdk_ptcatelh, smri_area_cdk_precnlh, smri_area_cdk_pclh, smri_area_cdk_rracatelh, smri_area_cdk_rrmdfrlh, smri_area_cdk_sufrlh, smri_area_cdk_supllh, smri_area_cdk_sutmlh, smri_area_cdk_smlh, smri_area_cdk_frpolelh, smri_area_cdk_tmpolelh, smri_area_cdk_trvtmlh, smri_area_cdk_insulalh, smri_area_cdk_banksstsrh, smri_area_cdk_cdacaterh, smri_area_cdk_cdmdfrrh, smri_area_cdk_cuneusrh, smri_area_cdk_ehinalrh, smri_area_cdk_fusiformrh, smri_area_cdk_ifplrh, smri_area_cdk_iftmrh, smri_area_cdk_ihcaterh, smri_area_cdk_loccrh, smri_area_cdk_lobfrrh, smri_area_cdk_lingualrh, smri_area_cdk_mobfrrh, smri_area_cdk_mdtmrh, smri_area_cdk_parahpalrh, smri_area_cdk_paracnrh, smri_area_cdk_parsopcrh, smri_area_cdk_parsobisrh, smri_area_cdk_parstgrisrh, smri_area_cdk_periccrh, smri_area_cdk_postcnrh, smri_area_cdk_ptcaterh, smri_area_cdk_precnrh, smri_area_cdk_pcrh, smri_area_cdk_rracaterh, smri_area_cdk_rrmdfrrh, smri_area_cdk_sufrrh, smri_area_cdk_suplrh, smri_area_cdk_sutmrh, smri_area_cdk_smrh, smri_area_cdk_frpolerh, smri_area_cdk_tmpolerh, smri_area_cdk_trvtmrh, smri_area_cdk_insularh, smri_area_cdk_totallh, smri_area_cdk_totalrh, smri_area_cdk_total, smri_vol_cdk_banksstslh, smri_vol_cdk_cdacatelh, smri_vol_cdk_cdmdfrlh, smri_vol_cdk_cuneuslh, smri_vol_cdk_ehinallh, smri_vol_cdk_fusiformlh, smri_vol_cdk_ifpllh, smri_vol_cdk_iftmlh, smri_vol_cdk_ihcatelh, smri_vol_cdk_locclh, smri_vol_cdk_lobfrlh, smri_vol_cdk_linguallh, smri_vol_cdk_mobfrlh, smri_vol_cdk_mdtmlh, smri_vol_cdk_parahpallh, smri_vol_cdk_paracnlh, smri_vol_cdk_parsopclh, smri_vol_cdk_parsobislh, smri_vol_cdk_parstgrislh, smri_vol_cdk_pericclh, smri_vol_cdk_postcnlh, smri_vol_cdk_ptcatelh, smri_vol_cdk_precnlh, smri_vol_cdk_pclh, smri_vol_cdk_rracatelh, smri_vol_cdk_rrmdfrlh, smri_vol_cdk_sufrlh, smri_vol_cdk_supllh, smri_vol_cdk_sutmlh, smri_vol_cdk_smlh, smri_vol_cdk_frpolelh, smri_vol_cdk_tmpolelh, smri_vol_cdk_trvtmlh, smri_vol_cdk_insulalh, smri_vol_cdk_banksstsrh, smri_vol_cdk_cdacaterh, smri_vol_cdk_cdmdfrrh, smri_vol_cdk_cuneusrh, smri_vol_cdk_ehinalrh, smri_vol_cdk_fusiformrh, smri_vol_cdk_ifplrh, smri_vol_cdk_iftmrh, smri_vol_cdk_ihcaterh, smri_vol_cdk_loccrh, smri_vol_cdk_lobfrrh, smri_vol_cdk_lingualrh, smri_vol_cdk_mobfrrh, smri_vol_cdk_mdtmrh, smri_vol_cdk_parahpalrh, smri_vol_cdk_paracnrh, smri_vol_cdk_parsopcrh, smri_vol_cdk_parsobisrh, smri_vol_cdk_parstgrisrh, smri_vol_cdk_periccrh, smri_vol_cdk_postcnrh, smri_vol_cdk_ptcaterh, smri_vol_cdk_precnrh, smri_vol_cdk_pcrh, smri_vol_cdk_rracaterh, smri_vol_cdk_rrmdfrrh, smri_vol_cdk_sufrrh, smri_vol_cdk_suplrh, smri_vol_cdk_sutmrh, smri_vol_cdk_smrh, smri_vol_cdk_frpolerh, smri_vol_cdk_tmpolerh, smri_vol_cdk_trvtmrh, smri_vol_cdk_insularh, smri_vol_cdk_totallh, smri_vol_cdk_totalrh, smri_vol_cdk_total, smri_vol_scs_cbwmatterlh, smri_vol_scs_ltventriclelh, smri_vol_scs_inflatventlh, smri_vol_scs_crbwmatterlh, smri_vol_scs_crbcortexlh, smri_vol_scs_tplh, smri_vol_scs_caudatelh, smri_vol_scs_putamenlh, smri_vol_scs_pallidumlh, smri_vol_scs_3rdventricle, smri_vol_scs_4thventricle, smri_vol_scs_bstem, smri_vol_scs_hpuslh, smri_vol_scs_amygdalalh, smri_vol_scs_csf, smri_vol_scs_aal, smri_vol_scs_vedclh, smri_vol_scs_cbwmatterrh, smri_vol_scs_ltventriclerh, smri_vol_scs_inflatventrh, smri_vol_scs_crbwmatterrh, smri_vol_scs_crbcortexrh, smri_vol_scs_tprh, smri_vol_scs_caudaterh, smri_vol_scs_putamenrh, smri_vol_scs_pallidumrh, smri_vol_scs_hpusrh, smri_vol_scs_amygdalarh, smri_vol_scs_aar, smri_vol_scs_vedcrh, smri_vol_scs_wmhint, smri_vol_scs_ccps, smri_vol_scs_ccmidps, smri_vol_scs_ccct, smri_vol_scs_ccmidat, smri_vol_scs_ccat, smri_vol_scs_wholeb, smri_vol_scs_latventricles, smri_vol_scs_allventricles, smri_vol_scs_intracranialv, smri_vol_scs_suprateialv, smri_vol_scs_subcorticalgv, smri_t1wcnt_cdk_banksstslh, smri_t1wcnt_cdk_cdacatelh, smri_t1wcnt_cdk_cdmdfrlh, smri_t1wcnt_cdk_cuneuslh, smri_t1wcnt_cdk_ehinallh, smri_t1wcnt_cdk_fusiformlh, smri_t1wcnt_cdk_ifpllh, smri_t1wcnt_cdk_iftmlh, smri_t1wcnt_cdk_ihcatelh, smri_t1wcnt_cdk_locclh, smri_t1wcnt_cdk_lobfrlh, smri_t1wcnt_cdk_linguallh, smri_t1wcnt_cdk_mobfrlh, smri_t1wcnt_cdk_mdtmlh, smri_t1wcnt_cdk_parahpallh, smri_t1wcnt_cdk_paracnlh, smri_t1wcnt_cdk_parsopclh, smri_t1wcnt_cdk_parsobislh, smri_t1wcnt_cdk_parstgrislh, smri_t1wcnt_cdk_pericclh, smri_t1wcnt_cdk_postcnlh, smri_t1wcnt_cdk_ptcatelh, smri_t1wcnt_cdk_precnlh, smri_t1wcnt_cdk_pclh, smri_t1wcnt_cdk_rracatelh, smri_t1wcnt_cdk_rrmdfrlh, smri_t1wcnt_cdk_sufrlh, smri_t1wcnt_cdk_supllh, smri_t1wcnt_cdk_sutmlh, smri_t1wcnt_cdk_smlh, smri_t1wcnt_cdk_frpolelh, smri_t1wcnt_cdk_tmpolelh, smri_t1wcnt_cdk_trvtmlh, smri_t1wcnt_cdk_insulalh, smri_t1wcnt_cdk_banksstsrh, smri_t1wcnt_cdk_cdacaterh, smri_t1wcnt_cdk_cdmdfrrh, smri_t1wcnt_cdk_cuneusrh, smri_t1wcnt_cdk_ehinalrh, smri_t1wcnt_cdk_fusiformrh, smri_t1wcnt_cdk_ifplrh, smri_t1wcnt_cdk_iftmrh, smri_t1wcnt_cdk_ihcaterh, smri_t1wcnt_cdk_loccrh, smri_t1wcnt_cdk_lobfrrh, smri_t1wcnt_cdk_lingualrh, smri_t1wcnt_cdk_mobfrrh, smri_t1wcnt_cdk_mdtmrh, smri_t1wcnt_cdk_parahpalrh, smri_t1wcnt_cdk_paracnrh, smri_t1wcnt_cdk_parsopcrh, smri_t1wcnt_cdk_parsobisrh, smri_t1wcnt_cdk_parstgrisrh, smri_t1wcnt_cdk_periccrh, smri_t1wcnt_cdk_postcnrh, smri_t1wcnt_cdk_ptcaterh, smri_t1wcnt_cdk_precnrh, smri_t1wcnt_cdk_pcrh, smri_t1wcnt_cdk_rracaterh, smri_t1wcnt_cdk_rrmdfrrh, smri_t1wcnt_cdk_sufrrh, smri_t1wcnt_cdk_suplrh, smri_t1wcnt_cdk_sutmrh, smri_t1wcnt_cdk_smrh, smri_t1wcnt_cdk_frpolerh, smri_t1wcnt_cdk_tmpolerh, smri_t1wcnt_cdk_trvtmrh, smri_t1wcnt_cdk_insularh, smri_t1wcnt_cdk_meanlh, smri_t1wcnt_cdk_meanrh, smri_t1wcnt_cdk_mean

**SI Table 3**. Showing list of tabulated DTI brain features (N = 576) extracted from the ABCD Study that was used for brain age prediction.

dmri_dtifa_fiberat_allfibers, dmri_dtifa_fiberat_allfiblh, dmri_dtifa_fiberat_allfcclh, dmri_dtifa_fiberat_allfccrh, dmri_dtifa_fiberat_allfibrh, dmri_dtifa_fiberat_atrlh, dmri_dtifa_fiberat_atrrh, dmri_dtifa_fiberat_cc, dmri_dtifa_fiberat_cgclh, dmri_dtifa_fiberat_cgcrh, dmri_dtifa_fiberat_cghlh, dmri_dtifa_fiberat_cghrh, dmri_dtifa_fiberat_cstlh, dmri_dtifa_fiberat_cstrh, dmri_dtifa_fiberat_fmaj, dmri_dtifa_fiberat_fmin, dmri_dtifa_fiberat_fscslh, dmri_dtifa_fiberat_fscsrh, dmri_dtifa_fiberat_fxcutlh, dmri_dtifa_fiberat_fxcutrh, dmri_dtifa_fiberat_fxlh, dmri_dtifa_fiberat_fxrh, dmri_dtifa_fiberat_ifolh, dmri_dtifa_fiberat_iforh, dmri_dtifa_fiberat_ifsfclh, dmri_dtifa_fiberat_ifsfcrh, dmri_dtifa_fiberat_ilflh, dmri_dtifa_fiberat_ilfrh, dmri_dtifa_fiberat_pscslh, dmri_dtifa_fiberat_pscsrh, dmri_dtifa_fiberat_pslflh, dmri_dtifa_fiberat_pslfrh, dmri_dtifa_fiberat_scslh, dmri_dtifa_fiberat_scsrh, dmri_dtifa_fiberat_sifclh, dmri_dtifa_fiberat_sifcrh, dmri_dtifa_fiberat_slflh, dmri_dtifa_fiberat_slfrh, dmri_dtifa_fiberat_tslflh, dmri_dtifa_fiberat_tslfrh, dmri_dtifa_fiberat_unclh, dmri_dtifa_fiberat_uncrh, dmri_dtimd_fiberat_allfibers, dmri_dtimd_fiberat_allfiblh, dmri_dtimd_fiberat_allfcclh, dmri_dtimd_fiberat_allfccrh, dmri_dtimd_fiberat_allfibrh, dmri_dtimd_fiberat_atrlh, dmri_dtimd_fiberat_atrrh, dmri_dtimd_fiberat_cc, dmri_dtimd_fiberat_cgclh, dmri_dtimd_fiberat_cgcrh, dmri_dtimd_fiberat_cghlh, dmri_dtimd_fiberat_cghrh, dmri_dtimd_fiberat_cstlh, dmri_dtimd_fiberat_cstrh, dmri_dtimd_fiberat_fmaj, dmri_dtimd_fiberat_fmin, dmri_dtimd_fiberat_fscslh, dmri_dtimd_fiberat_fscsrh, dmri_dtimd_fiberat_fxcutlh, dmri_dtimd_fiberat_fxcutrh, dmri_dtimd_fiberat_fxlh, dmri_dtimd_fiberat_fxrh, dmri_dtimd_fiberat_ifolh, dmri_dtimd_fiberat_iforh, dmri_dtimd_fiberat_ifsfclh, dmri_dtimd_fiberat_ifsfcrh, dmri_dtimd_fiberat_ilflh, dmri_dtimd_fiberat_ilfrh, dmri_dtimd_fiberat_pscslh, dmri_dtimd_fiberat_pscsrh, dmri_dtimd_fiberat_pslflh, dmri_dtimd_fiberat_pslfrh, dmri_dtimd_fiberat_scslh, dmri_dtimd_fiberat_scsrh, dmri_dtimd_fiberat_sifclh, dmri_dtimd_fiberat_sifcrh, dmri_dtimd_fiberat_slflh, dmri_dtimd_fiberat_slfrh, dmri_dtimd_fiberat_tslflh, dmri_dtimd_fiberat_tslfrh, dmri_dtimd_fiberat_unclh, dmri_dtimd_fiberat_uncrh, dmri_dtild_fiberat_allfibers, dmri_dtild_fiberat_allfiblh, dmri_dtild_fiberat_allfocclh, dmri_dtild_fiberat_allfccrh, dmri_dtild_fiberat_allfibrh, dmri_dtild_fiberat_atrlh, dmri_dtild_fiberat_atrrh, dmri_dtild_fiberat_cc, dmri_dtild_fiberat_cgclh, dmri_dtild_fiberat_cgcrh, dmri_dtild_fiberat_cghlh, dmri_dtild_fiberat_cghrh, dmri_dtild_fiberat_cstlh, dmri_dtild_fiberat_cstrh, dmri_dtild_fiberat_fmaj, dmri_dtild_fiberat_fmin, dmri_dtild_fiberat_fscslh, dmri_dtild_fiberat_fscsrh, dmri_dtild_fiberat_fxcutlh, dmri_dtild_fiberat_fxcutrh, dmri_dtild_fiberat_fxlh, dmri_dtild_fiberat_fxrh, dmri_dtild_fiberat_ifolh, dmri_dtild_fiberat_iforh, dmri_dtild_fiberat_ifsfclh, dmri_dtild_fiberat_ifsfcrh, dmri_dtild_fiberat_ilflh, dmri_dtild_fiberat_ilfrh, dmri_dtild_fiberat_pscslh, dmri_dtild_fiberat_pscsrh, dmri_dtild_fiberat_pslflh, dmri_dtild_fiberat_pslfrh, dmri_dtild_fiberat_scslh, dmri_dtild_fiberat_scsrh, dmri_dtild_fiberat_sifclh, dmri_dtild_fiberat_sifcrh, dmri_dtild_fiberat_slflh, dmri_dtild_fiberat_slfrh, dmri_dtild_fiberat_tslflh, dmri_dtild_fiberat_tslfrh, dmri_dtild_fiberat_unclh, dmri_dtild_fiberat_uncrh, dmri_dtitd_fiberat_allfibers, dmri_dtitd_fiberat_allfiblh, dmri_dtitd_fiberat_allfcclh, dmri_dtitd_fiberat_allfccrh, dmri_dtitd_fiberat_allfibrh, dmri_dtitd_fiberat_atrlh, dmri_dtitd_fiberat_atrrh, dmri_dtitd_fiberat_cc, dmri_dtitd_fiberat_cgclh, dmri_dtitd_fiberat_cgcrh, dmri_dtitd_fiberat_cghlh, dmri_dtitd_fiberat_cghrh, dmri_dtitd_fiberat_cstlh, dmri_dtitd_fiberat_cstrh, dmri_dtitd_fiberat_fmaj, dmri_dtitd_fiberat_fmin, dmri_dtitd_fiberat_fscslh, dmri_dtitd_fiberat_fscsrh, dmri_dtitd_fiberat_fxcutlh, dmri_dtitd_fiberat_fxcutrh, dmri_dtitd_fiberat_fxlh, dmri_dtitd_fiberat_fxrh, dmri_dtitd_fiberat_ifolh, dmri_dtitd_fiberat_iforh, dmri_dtitd_fiberat_ifsfclh, dmri_dtitd_fiberat_ifsfcrh, dmri_dtitd_fiberat_ilflh, dmri_dtitd_fiberat_ilfrh, dmri_dtitd_fiberat_pscslh, dmri_dtitd_fiberat_pscsrh, dmri_dtitd_fiberat_pslflh, dmri_dtitd_fiberat_pslfrh, dmri_dtitd_fiberat_scslh, dmri_dtitd_fiberat_scsrh, dmri_dtitd_fiberat_sifclh, dmri_dtitd_fiberat_sifcrh, dmri_dtitd_fiberat_slflh, dmri_dtitd_fiberat_slfrh, dmri_dtitd_fiberat_tslflh, dmri_dtitd_fiberat_tslfrh, dmri_dtitd_fiberat_unclh, dmri_dtitd_fiberat_uncrh, dmri_dtifa_scts_3rdvt, dmri_dtifa_scts_4thvt, dmri_dtifa_scts_accarealh, dmri_dtifa_scts_accarearh, dmri_dtifa_scts_amygdalalh, dmri_dtifa_scts_amygdalarh, dmri_dtifa_scts_brainstem, dmri_dtifa_scts_caudatelh, dmri_dtifa_scts_caudaterh, dmri_dtifa_scts_cblcortexlh, dmri_dtifa_scts_cblcortexrh, dmri_dtifa_scts_cblwmlh, dmri_dtifa_scts_cblwmrh, dmri_dtifa_scts_crwmlh, dmri_dtifa_scts_crwmrh, dmri_dtifa_scts_csf, dmri_dtifa_scts_hpuslh, dmri_dtifa_scts_hpusrh, dmri_dtifa_scts_inflatventlh, dmri_dtifa_scts_inflatventrh, dmri_dtifa_scts_lvtlh, dmri_dtifa_scts_lvtrh, dmri_dtifa_scts_pdlh, dmri_dtifa_scts_pdrh, dmri_dtifa_scts_putamenlh, dmri_dtifa_scts_putamenrh, dmri_dtifa_scts_thplh, dmri_dtifa_scts_thprh, dmri_dtifa_scts_ventraldclh, dmri_dtifa_scts_ventraldcrh, dmri_dtimd_scts_3rdvt, dmri_dtimd_scts_4thvt, dmri_dtimd_scts_accarealh, dmri_dtimd_scts_accarearh, dmri_dtimd_scts_amygdalalh, dmri_dtimd_scts_amygdalarh, dmri_dtimd_scts_brainstem, dmri_dtimd_scts_caudatelh, dmri_dtimd_scts_caudaterh, dmri_dtimd_scts_cblcortexlh, dmri_dtimd_scts_cblcortexrh, dmri_dtimd_scts_cblwmlh, dmri_dtimd_scts_cblwmrh, dmri_dtimd_scts_crwmlh, dmri_dtimd_scts_crwmrh, dmri_dtimd_scts_csf, dmri_dtimd_scts_hpuslh, dmri_dtimd_scts_hpusrh, dmri_dtimd_scts_inflatventlh, dmri_dtimd_scts_inflatventrh, dmri_dtimd_scts_lvtlh, dmri_dtimd_scts_lvtrh, dmri_dtimd_scts_pdlh, dmri_dtimd_scts_pdrh, dmri_dtimd_scts_putamenlh, dmri_dtimd_scts_putamenrh, dmri_dtimd_scts_thplh, dmri_dtimd_scts_thprh, dmri_dtimd_scts_ventraldclh, dmri_dtimd_scts_ventraldcrh, dmri_dtild_scts_3rdvt, dmri_dtild_scts_4thvt, dmri_dtild_scts_accarealh, dmri_dtild_scts_accarearh, dmri_dtild_scts_amygdalalh, dmri_dtild_scts_amygdalarh, dmri_dtild_scts_brainstem, dmri_dtild_scts_caudatelh, dmri_dtild_scts_caudaterh, dmri_dtild_scts_cblcortexlh, dmri_dtild_scts_cblcortexrh, dmri_dtild_scts_cblwmlh, dmri_dtild_scts_cblwmrh, dmri_dtild_scts_crwmlh, dmri_dtild_scts_crwmrh, dmri_dtild_scts_csf, dmri_dtild_scts_hpuslh, dmri_dtild_scts_hpusrh, dmri_dtild_scts_inflatventlh, dmri_dtild_scts_inflatventrh, dmri_dtild_scts_lvtlh, dmri_dtild_scts_lvtrh, dmri_dtild_scts_pdlh, dmri_dtild_scts_pdrh, dmri_dtild_scts_putamenlh, dmri_dtild_scts_putamenrh, dmri_dtild_scts_thplh, dmri_dtild_scts_thprh, dmri_dtild_scts_ventraldclh, dmri_dtild_scts_ventraldcrh, dmri_dtitd_scts_3rdvt, dmri_dtitd_scts_4thvt, dmri_dtitd_scts_accarealh, dmri_dtitd_scts_accarearh, dmri_dtitd_scts_amygdalalh, dmri_dtitd_scts_amygdalarh, dmri_dtitd_scts_brainstem, dmri_dtitd_scts_caudatelh, dmri_dtitd_scts_caudaterh, dmri_dtitd_scts_cblcortexlh, dmri_dtitd_scts_cblcortexrh, dmri_dtitd_scts_cblwmlh, dmri_dtitd_scts_cblwmrh, dmri_dtitd_scts_crwmlh, dmri_dtitd_scts_crwmrh, dmri_dtitd_scts_csf, dmri_dtitd_scts_hpuslh, dmri_dtitd_scts_hpusrh, dmri_dtitd_scts_inflatventlh, dmri_dtitd_scts_inflatventrh, dmri_dtitd_scts_lvtlh, dmri_dtitd_scts_lvtrh, dmri_dtitd_scts_pdlh, dmri_dtitd_scts_pdrh, dmri_dtitd_scts_putamenlh, dmri_dtitd_scts_putamenrh, dmri_dtitd_scts_thplh, dmri_dtitd_scts_thprh, dmri_dtitd_scts_ventraldclh, dmri_dtitd_scts_ventraldcrh, dmdtifp1_38, dmdtifp1_42, dmdtifp1_40, dmdtifp1_39, dmdtifp1_41, dmdtifp1_10, dmdtifp1_9, dmdtifp1_19, dmdtifp1_4, dmdtifp1_3, dmdtifp1_6, dmdtifp1_5, dmdtifp1_8, dmdtifp1_7, dmdtifp1_17, dmdtifp1_18, dmdtifp1_29, dmdtifp1_28, dmdtifp1_37, dmdtifp1_36, dmdtifp1_2, dmdtifp1_1, dmdtifp1_16, dmdtifp1_15, dmdtifp1_35, dmdtifp1_34, dmdtifp1_14, dmdtifp1_13, dmdtifp1_31, dmdtifp1_30, dmdtifp1_25, dmdtifp1_24, dmdtifp1_27, dmdtifp1_26, dmdtifp1_33, dmdtifp1_32, dmdtifp1_21, dmdtifp1_20, dmdtifp1_23, dmdtifp1_22, dmdtifp1_12, dmdtifp1_11, dmdtifp1_80, dmdtifp1_84, dmdtifp1_82, dmdtifp1_81, dmdtifp1_83, dmdtifp1_52, dmdtifp1_51, dmdtifp1_61, dmdtifp1_46, dmdtifp1_45, dmdtifp1_48, dmdtifp1_47, dmdtifp1_50, dmdtifp1_49, dmdtifp1_59, dmdtifp1_60, dmdtifp1_71, dmdtifp1_70, dmdtifp1_79, dmdtifp1_78, dmdtifp1_44, dmdtifp1_43, dmdtifp1_58, dmdtifp1_57, dmdtifp1_77, dmdtifp1_76, dmdtifp1_56, dmdtifp1_55, dmdtifp1_73, dmdtifp1_72, dmdtifp1_67, dmdtifp1_66, dmdtifp1_69, dmdtifp1_68, dmdtifp1_75, dmdtifp1_74, dmdtifp1_63, dmdtifp1_62, dmdtifp1_65, dmdtifp1_64, dmdtifp1_54, dmdtifp1_53, dmdtifp1_122, dmdtifp1_126, dmdtifp1_124, dmdtifp1_123, dmdtifp1_125, dmdtifp1_94, dmdtifp1_93, dmdtifp1_103, dmdtifp1_88, dmdtifp1_87, dmdtifp1_90, dmdtifp1_89, dmdtifp1_92, dmdtifp1_91, dmdtifp1_101, dmdtifp1_102, dmdtifp1_113, dmdtifp1_112, dmdtifp1_121, dmdtifp1_120, dmdtifp1_86, dmdtifp1_85, dmdtifp1_100, dmdtifp1_99, dmdtifp1_119, dmdtifp1_118, dmdtifp1_98, dmdtifp1_97, dmdtifp1_115, dmdtifp1_114, dmdtifp1_109, dmdtifp1_108, dmdtifp1_111, dmdtifp1_110, dmdtifp1_117, dmdtifp1_116, dmdtifp1_105, dmdtifp1_104, dmdtifp1_107, dmdtifp1_106, dmdtifp1_96, dmdtifp1_95, dmdtifp1_164, dmdtifp1_168, dmdtifp1_166, dmdtifp1_165, dmdtifp1_167, dmdtifp1_136, dmdtifp1_135, dmdtifp1_145, dmdtifp1_130, dmdtifp1_129, dmdtifp1_132, dmdtifp1_131, dmdtifp1_134, dmdtifp1_133, dmdtifp1_143, dmdtifp1_144, dmdtifp1_155, dmdtifp1_154, dmdtifp1_163, dmdtifp1_162, dmdtifp1_128, dmdtifp1_127, dmdtifp1_142, dmdtifp1_141, dmdtifp1_161, dmdtifp1_160, dmdtifp1_140, dmdtifp1_139, dmdtifp1_157, dmdtifp1_156, dmdtifp1_151, dmdtifp1_150, dmdtifp1_153, dmdtifp1_152, dmdtifp1_159, dmdtifp1_158, dmdtifp1_147, dmdtifp1_146, dmdtifp1_149, dmdtifp1_148, dmdtifp1_138, dmdtifp1_137, dmdtifp1_220, dmdtifp1_221, dmdtifp1_226, dmdtifp1_239, dmdtifp1_224, dmdtifp1_238, dmdtifp1_222, dmdtifp1_217, dmdtifp1_234, dmdtifp1_215, dmdtifp1_232, dmdtifp1_214, dmdtifp1_231, dmdtifp1_211, dmdtifp1_228, dmdtifp1_225, dmdtifp1_223, dmdtifp1_237, dmdtifp1_213, dmdtifp1_230, dmdtifp1_212, dmdtifp1_229, dmdtifp1_219, dmdtifp1_236, dmdtifp1_218, dmdtifp1_235, dmdtifp1_216, dmdtifp1_233, dmdtifp1_227, dmdtifp1_240, dmdtifp1_250, dmdtifp1_251, dmdtifp1_256, dmdtifp1_269, dmdtifp1_254, dmdtifp1_268, dmdtifp1_252, dmdtifp1_247, dmdtifp1_264, dmdtifp1_245, dmdtifp1_262, dmdtifp1_244, dmdtifp1_261, dmdtifp1_241, dmdtifp1_258, dmdtifp1_255, dmdtifp1_253, dmdtifp1_267, dmdtifp1_243, dmdtifp1_260, dmdtifp1_242, dmdtifp1_259, dmdtifp1_249, dmdtifp1_266, dmdtifp1_248, dmdtifp1_265, dmdtifp1_246, dmdtifp1_263, dmdtifp1_257, dmdtifp1_270, dmdtifp1_280, dmdtifp1_281, dmdtifp1_286, dmdtifp1_299, dmdtifp1_284, dmdtifp1_298, dmdtifp1_282, dmdtifp1_277, dmdtifp1_294, dmdtifp1_275, dmdtifp1_292, dmdtifp1_274, dmdtifp1_291, dmdtifp1_271, dmdtifp1_288, dmdtifp1_285, dmdtifp1_283, dmdtifp1_297, dmdtifp1_273, dmdtifp1_290, dmdtifp1_272, dmdtifp1_289, dmdtifp1_279, dmdtifp1_296, dmdtifp1_278, dmdtifp1_295, dmdtifp1_276, dmdtifp1_293, dmdtifp1_287, dmdtifp1_300, dmdtifp1_310, dmdtifp1_311, dmdtifp1_316, dmdtifp1_329, dmdtifp1_314, dmdtifp1_328, dmdtifp1_312, dmdtifp1_307, dmdtifp1_324, dmdtifp1_305, dmdtifp1_322, dmdtifp1_304, dmdtifp1_321, dmdtifp1_301, dmdtifp1_318, dmdtifp1_315, dmdtifp1_313, dmdtifp1_327, dmdtifp1_303, dmdtifp1_320, dmdtifp1_302, dmdtifp1_319, dmdtifp1_309, dmdtifp1_326, dmdtifp1_308, dmdtifp1_325, dmdtifp1_306, dmdtifp1_323, dmdtifp1_317, dmdtifp1_330

**SI Table 4**. Showing list of tabulated rs-fMRI brain features (N = 416) extracted from the ABCD Study that was used for brain age prediction.

rsfmri_c_ngd_ad_ngd_ad, rsfmri_c_ngd_ad_ngd_cgc, rsfmri_c_ngd_ad_ngd_ca, rsfmri_c_ngd_ad_ngd_dt, rsfmri_c_ngd_ad_ngd_dla, rsfmri_c_ngd_ad_ngd_fo, rsfmri_c_ngd_ad_ngd_n, rsfmri_c_ngd_ad_ngd_rspltp, rsfmri_c_ngd_ad_ngd_sa, rsfmri_c_ngd_ad_ngd_smh, rsfmri_c_ngd_ad_ngd_smm, rsfmri_c_ngd_ad_ngd_vta, rsfmri_c_ngd_ad_ngd_vs, rsfmri_c_ngd_cgc_ngd_ad, rsfmri_c_ngd_cgc_ngd_cgc, rsfmri_c_ngd_cgc_ngd_ca, rsfmri_c_ngd_cgc_ngd_dt, rsfmri_c_ngd_cgc_ngd_dla, rsfmri_c_ngd_cgc_ngd_fo, rsfmri_c_ngd_cgc_ngd_n, rsfmri_c_ngd_cgc_ngd_rspltp, rsfmri_c_ngd_cgc_ngd_sa, rsfmri_c_ngd_cgc_ngd_smh, rsfmri_c_ngd_cgc_ngd_smm, rsfmri_c_ngd_cgc_ngd_vta, rsfmri_c_ngd_cgc_ngd_vs, rsfmri_c_ngd_ca_ngd_ad, rsfmri_c_ngd_ca_ngd_cgc, rsfmri_c_ngd_ca_ngd_ca, rsfmri_c_ngd_ca_ngd_dt, rsfmri_c_ngd_ca_ngd_dla, rsfmri_c_ngd_ca_ngd_fo, rsfmri_c_ngd_ca_ngd_n, rsfmri_c_ngd_ca_ngd_rspltp, rsfmri_c_ngd_ca_ngd_sa, rsfmri_c_ngd_ca_ngd_smh, rsfmri_c_ngd_ca_ngd_smm, rsfmri_c_ngd_ca_ngd_vta, rsfmri_c_ngd_ca_ngd_vs, rsfmri_c_ngd_dt_ngd_ad, rsfmri_c_ngd_dt_ngd_cgc, rsfmri_c_ngd_dt_ngd_ca, rsfmri_c_ngd_dt_ngd_dt, rsfmri_c_ngd_dt_ngd_dla, rsfmri_c_ngd_dt_ngd_fo, rsfmri_c_ngd_dt_ngd_n, rsfmri_c_ngd_dt_ngd_rspltp, rsfmri_c_ngd_dt_ngd_sa, rsfmri_c_ngd_dt_ngd_smh, rsfmri_c_ngd_dt_ngd_smm, rsfmri_c_ngd_dt_ngd_vta, rsfmri_c_ngd_dt_ngd_vs, rsfmri_c_ngd_dla_ngd_ad, rsfmri_c_ngd_dla_ngd_cgc, rsfmri_c_ngd_dla_ngd_ca, rsfmri_c_ngd_dla_ngd_dt, rsfmri_c_ngd_dla_ngd_dla, rsfmri_c_ngd_dla_ngd_fo, rsfmri_c_ngd_dla_ngd_n, rsfmri_c_ngd_dla_ngd_rspltp, rsfmri_c_ngd_dla_ngd_sa, rsfmri_c_ngd_dla_ngd_smh, rsfmri_c_ngd_dla_ngd_smm, rsfmri_c_ngd_dla_ngd_vta, rsfmri_c_ngd_dla_ngd_vs, rsfmri_c_ngd_fo_ngd_ad, rsfmri_c_ngd_fo_ngd_cgc, rsfmri_c_ngd_fo_ngd_ca, rsfmri_c_ngd_fo_ngd_dt, rsfmri_c_ngd_fo_ngd_dla, rsfmri_c_ngd_fo_ngd_fo, rsfmri_c_ngd_fo_ngd_n, rsfmri_c_ngd_fo_ngd_rspltp, rsfmri_c_ngd_fo_ngd_sa, rsfmri_c_ngd_fo_ngd_smh, rsfmri_c_ngd_fo_ngd_smm, rsfmri_c_ngd_fo_ngd_vta, rsfmri_c_ngd_fo_ngd_vs, rsfmri_c_ngd_n_ngd_ad, rsfmri_c_ngd_n_ngd_cgc, rsfmri_c_ngd_n_ngd_ca, rsfmri_c_ngd_n_ngd_dt, rsfmri_c_ngd_n_ngd_dla, rsfmri_c_ngd_n_ngd_fo, rsfmri_c_ngd_n_ngd_n, rsfmri_c_ngd_n_ngd_rspltp, rsfmri_c_ngd_n_ngd_sa, rsfmri_c_ngd_n_ngd_smh, rsfmri_c_ngd_n_ngd_smm, rsfmri_c_ngd_n_ngd_vta, rsfmri_c_ngd_n_ngd_vs, rsfmri_c_ngd_rspltp_ngd_ad, rsfmri_c_ngd_rspltp_ngd_cgc, rsfmri_c_ngd_rspltp_ngd_ca, rsfmri_c_ngd_rspltp_ngd_dt, rsfmri_c_ngd_rspltp_ngd_dla, rsfmri_c_ngd_rspltp_ngd_fo, rsfmri_c_ngd_rspltp_ngd_n, rsfmri_c_ngd_rspltp_ngd_rspltp, rsfmri_c_ngd_rspltp_ngd_sa, rsfmri_c_ngd_rspltp_ngd_smh, rsfmri_c_ngd_rspltp_ngd_smm, rsfmri_c_ngd_rspltp_ngd_vta, rsfmri_c_ngd_rspltp_ngd_vs, rsfmri_c_ngd_sa_ngd_ad, rsfmri_c_ngd_sa_ngd_cgc, rsfmri_c_ngd_sa_ngd_ca, rsfmri_c_ngd_sa_ngd_dt, rsfmri_c_ngd_sa_ngd_dla, rsfmri_c_ngd_sa_ngd_fo, rsfmri_c_ngd_sa_ngd_n, rsfmri_c_ngd_sa_ngd_rspltp, rsfmri_c_ngd_sa_ngd_sa, rsfmri_c_ngd_sa_ngd_smh, rsfmri_c_ngd_sa_ngd_smm, rsfmri_c_ngd_sa_ngd_vta, rsfmri_c_ngd_sa_ngd_vs, rsfmri_c_ngd_smh_ngd_ad, rsfmri_c_ngd_smh_ngd_cgc, rsfmri_c_ngd_smh_ngd_ca, rsfmri_c_ngd_smh_ngd_dt, rsfmri_c_ngd_smh_ngd_dla, rsfmri_c_ngd_smh_ngd_fo, rsfmri_c_ngd_smh_ngd_n, rsfmri_c_ngd_smh_ngd_rspltp, rsfmri_c_ngd_smh_ngd_sa, rsfmri_c_ngd_smh_ngd_smh, rsfmri_c_ngd_smh_ngd_smm, rsfmri_c_ngd_smh_ngd_vta, rsfmri_c_ngd_smh_ngd_vs, rsfmri_c_ngd_smm_ngd_ad, rsfmri_c_ngd_smm_ngd_cgc, rsfmri_c_ngd_smm_ngd_ca, rsfmri_c_ngd_smm_ngd_dt, rsfmri_c_ngd_smm_ngd_dla, rsfmri_c_ngd_smm_ngd_fo, rsfmri_c_ngd_smm_ngd_n, rsfmri_c_ngd_smm_ngd_rspltp, rsfmri_c_ngd_smm_ngd_sa, rsfmri_c_ngd_smm_ngd_smh, rsfmri_c_ngd_smm_ngd_smm, rsfmri_c_ngd_smm_ngd_vta, rsfmri_c_ngd_smm_ngd_vs, rsfmri_c_ngd_vta_ngd_ad, rsfmri_c_ngd_vta_ngd_cgc, rsfmri_c_ngd_vta_ngd_ca, rsfmri_c_ngd_vta_ngd_dt, rsfmri_c_ngd_vta_ngd_dla, rsfmri_c_ngd_vta_ngd_fo, rsfmri_c_ngd_vta_ngd_n, rsfmri_c_ngd_vta_ngd_rspltp, rsfmri_c_ngd_vta_ngd_sa, rsfmri_c_ngd_vta_ngd_smh, rsfmri_c_ngd_vta_ngd_smm, rsfmri_c_ngd_vta_ngd_vta, rsfmri_c_ngd_vta_ngd_vs, rsfmri_c_ngd_vs_ngd_ad, rsfmri_c_ngd_vs_ngd_cgc, rsfmri_c_ngd_vs_ngd_ca, rsfmri_c_ngd_vs_ngd_dt, rsfmri_c_ngd_vs_ngd_dla, rsfmri_c_ngd_vs_ngd_fo, rsfmri_c_ngd_vs_ngd_n, rsfmri_c_ngd_vs_ngd_rspltp, rsfmri_c_ngd_vs_ngd_sa, rsfmri_c_ngd_vs_ngd_smh, rsfmri_c_ngd_vs_ngd_smm, rsfmri_c_ngd_vs_ngd_vta, rsfmri_c_ngd_vs_ngd_vs, rsfmri_cor_ngd_au_scs_aalh, rsfmri_cor_ngd_au_scs_aarh, rsfmri_cor_ngd_au_scs_aglh, rsfmri_cor_ngd_au_scs_agrh, rsfmri_cor_ngd_au_scs_bs, rsfmri_cor_ngd_au_scs_cdelh, rsfmri_cor_ngd_au_scs_cderh, rsfmri_cor_ngd_au_scs_crcxlh, rsfmri_cor_ngd_au_scs_crcxrh, rsfmri_cor_ngd_au_scs_hplh, rsfmri_cor_ngd_au_scs_hprh, rsfmri_cor_ngd_au_scs_pllh, rsfmri_cor_ngd_au_scs_plrh, rsfmri_cor_ngd_au_scs_ptlh, rsfmri_cor_ngd_au_scs_ptrh, rsfmri_cor_ngd_au_scs_thplh, rsfmri_cor_ngd_au_scs_thprh, rsfmri_cor_ngd_au_scs_vtdclh, rsfmri_cor_ngd_au_scs_vtdcrh, rsfmri_cor_ngd_cerc_scs_aalh, rsfmri_cor_ngd_cerc_scs_aarh, rsfmri_cor_ngd_cerc_scs_aglh, rsfmri_cor_ngd_cerc_scs_agrh, rsfmri_cor_ngd_cerc_scs_bs, rsfmri_cor_ngd_cerc_scs_cdelh, rsfmri_cor_ngd_cerc_scs_cderh, rsfmri_cor_ngd_cerc_scs_crcxlh, rsfmri_cor_ngd_cerc_scs_crcxrh, rsfmri_cor_ngd_cerc_scs_hplh, rsfmri_cor_ngd_cerc_scs_hprh, rsfmri_cor_ngd_cerc_scs_pllh, rsfmri_cor_ngd_cerc_scs_plrh, rsfmri_cor_ngd_cerc_scs_ptlh, rsfmri_cor_ngd_cerc_scs_ptrh, rsfmri_cor_ngd_cerc_scs_thplh, rsfmri_cor_ngd_cerc_scs_thprh, rsfmri_cor_ngd_cerc_scs_vtdclh, rsfmri_cor_ngd_cerc_scs_vtdcrh, rsfmri_cor_ngd_copa_scs_aalh, rsfmri_cor_ngd_copa_scs_aarh, rsfmri_cor_ngd_copa_scs_aglh, rsfmri_cor_ngd_copa_scs_agrh, rsfmri_cor_ngd_copa_scs_bs, rsfmri_cor_ngd_copa_scs_cdelh, rsfmri_cor_ngd_copa_scs_cderh, rsfmri_cor_ngd_copa_scs_crcxlh, rsfmri_cor_ngd_copa_scs_crcxrh, rsfmri_cor_ngd_copa_scs_hplh, rsfmri_cor_ngd_copa_scs_hprh, rsfmri_cor_ngd_copa_scs_pllh, rsfmri_cor_ngd_copa_scs_plrh, rsfmri_cor_ngd_copa_scs_ptlh, rsfmri_cor_ngd_copa_scs_ptrh, rsfmri_cor_ngd_copa_scs_thplh, rsfmri_cor_ngd_copa_scs_thprh, rsfmri_cor_ngd_copa_scs_vtdclh, rsfmri_cor_ngd_copa_scs_vtdcrh, rsfmri_cor_ngd_df_scs_aalh, rsfmri_cor_ngd_df_scs_aarh, rsfmri_cor_ngd_df_scs_aglh, rsfmri_cor_ngd_df_scs_agrh, rsfmri_cor_ngd_df_scs_bs, rsfmri_cor_ngd_df_scs_cdelh, rsfmri_cor_ngd_df_scs_cderh, rsfmri_cor_ngd_df_scs_crcxlh, rsfmri_cor_ngd_df_scs_crcxrh, rsfmri_cor_ngd_df_scs_hplh, rsfmri_cor_ngd_df_scs_hprh, rsfmri_cor_ngd_df_scs_pllh, rsfmri_cor_ngd_df_scs_plrh, rsfmri_cor_ngd_df_scs_ptlh, rsfmri_cor_ngd_df_scs_ptrh, rsfmri_cor_ngd_df_scs_thplh, rsfmri_cor_ngd_df_scs_thprh, rsfmri_cor_ngd_df_scs_vtdclh, rsfmri_cor_ngd_df_scs_vtdcrh, rsfmri_cor_ngd_dsa_scs_aalh, rsfmri_cor_ngd_dsa_scs_aarh, rsfmri_cor_ngd_dsa_scs_aglh, rsfmri_cor_ngd_dsa_scs_agrh, rsfmri_cor_ngd_dsa_scs_bs, rsfmri_cor_ngd_dsa_scs_cdelh, rsfmri_cor_ngd_dsa_scs_cderh, rsfmri_cor_ngd_dsa_scs_crcxlh, rsfmri_cor_ngd_dsa_scs_crcxrh, rsfmri_cor_ngd_dsa_scs_hplh, rsfmri_cor_ngd_dsa_scs_hprh, rsfmri_cor_ngd_dsa_scs_pllh, rsfmri_cor_ngd_dsa_scs_plrh, rsfmri_cor_ngd_dsa_scs_ptlh, rsfmri_cor_ngd_dsa_scs_ptrh, rsfmri_cor_ngd_dsa_scs_thplh, rsfmri_cor_ngd_dsa_scs_thprh, rsfmri_cor_ngd_dsa_scs_vtdclh, rsfmri_cor_ngd_dsa_scs_vtdcrh, rsfmri_cor_ngd_fopa_scs_aalh, rsfmri_cor_ngd_fopa_scs_aarh, rsfmri_cor_ngd_fopa_scs_aglh, rsfmri_cor_ngd_fopa_scs_agrh, rsfmri_cor_ngd_fopa_scs_bs, rsfmri_cor_ngd_fopa_scs_cdelh, rsfmri_cor_ngd_fopa_scs_cderh, rsfmri_cor_ngd_fopa_scs_crcxlh, rsfmri_cor_ngd_fopa_scs_crcxrh, rsfmri_cor_ngd_fopa_scs_hplh, rsfmri_cor_ngd_fopa_scs_hprh, rsfmri_cor_ngd_fopa_scs_pllh, rsfmri_cor_ngd_fopa_scs_plrh, rsfmri_cor_ngd_fopa_scs_ptlh, rsfmri_cor_ngd_fopa_scs_ptrh, rsfmri_cor_ngd_fopa_scs_thplh, rsfmri_cor_ngd_fopa_scs_thprh, rsfmri_cor_ngd_fopa_scs_vtdclh, rsfmri_cor_ngd_fopa_scs_vtdcrh, rsfmri_cor_ngd_none_scs_aalh, rsfmri_cor_ngd_none_scs_aarh, rsfmri_cor_ngd_none_scs_aglh, rsfmri_cor_ngd_none_scs_agrh, rsfmri_cor_ngd_none_scs_bs, rsfmri_cor_ngd_none_scs_cdelh, rsfmri_cor_ngd_none_scs_cderh, rsfmri_cor_ngd_none_scs_crcxlh, rsfmri_cor_ngd_none_scs_crcxrh, rsfmri_cor_ngd_none_scs_hplh, rsfmri_cor_ngd_none_scs_hprh, rsfmri_cor_ngd_none_scs_pllh, rsfmri_cor_ngd_none_scs_plrh, rsfmri_cor_ngd_none_scs_ptlh, rsfmri_cor_ngd_none_scs_ptrh, rsfmri_cor_ngd_none_scs_thplh, rsfmri_cor_ngd_none_scs_thprh, rsfmri_cor_ngd_none_scs_vtdclh, rsfmri_cor_ngd_none_scs_vtdcrh, rsfmri_cor_ngd_rst_scs_aalh, rsfmri_cor_ngd_rst_scs_aarh, rsfmri_cor_ngd_rst_scs_aglh, rsfmri_cor_ngd_rst_scs_agrh, rsfmri_cor_ngd_rst_scs_bs, rsfmri_cor_ngd_rst_scs_cdelh, rsfmri_cor_ngd_rst_scs_cderh, rsfmri_cor_ngd_rst_scs_crcxlh, rsfmri_cor_ngd_rst_scs_crcxrh, rsfmri_cor_ngd_rst_scs_hplh, rsfmri_cor_ngd_rst_scs_hprh, rsfmri_cor_ngd_rst_scs_pllh, rsfmri_cor_ngd_rst_scs_plrh, rsfmri_cor_ngd_rst_scs_ptlh, rsfmri_cor_ngd_rst_scs_ptrh, rsfmri_cor_ngd_rst_scs_thplh, rsfmri_cor_ngd_rst_scs_thprh, rsfmri_cor_ngd_rst_scs_vtdclh, rsfmri_cor_ngd_rst_scs_vtdcrh, rsfmri_cor_ngd_sa_scs_aalh, rsfmri_cor_ngd_sa_scs_aarh, rsfmri_cor_ngd_sa_scs_aglh, rsfmri_cor_ngd_sa_scs_agrh, rsfmri_cor_ngd_sa_scs_bs, rsfmri_cor_ngd_sa_scs_cdelh, rsfmri_cor_ngd_sa_scs_cderh, rsfmri_cor_ngd_sa_scs_crcxlh, rsfmri_cor_ngd_sa_scs_crcxrh, rsfmri_cor_ngd_sa_scs_hplh, rsfmri_cor_ngd_sa_scs_hprh, rsfmri_cor_ngd_sa_scs_pllh, rsfmri_cor_ngd_sa_scs_plrh, rsfmri_cor_ngd_sa_scs_ptlh, rsfmri_cor_ngd_sa_scs_ptrh, rsfmri_cor_ngd_sa_scs_thplh, rsfmri_cor_ngd_sa_scs_thprh, rsfmri_cor_ngd_sa_scs_vtdclh, rsfmri_cor_ngd_sa_scs_vtdcrh, rsfmri_cor_ngd_smh_scs_aalh, rsfmri_cor_ngd_smh_scs_aarh, rsfmri_cor_ngd_smh_scs_aglh, rsfmri_cor_ngd_smh_scs_agrh, rsfmri_cor_ngd_smh_scs_bs, rsfmri_cor_ngd_smh_scs_cdelh, rsfmri_cor_ngd_smh_scs_cderh, rsfmri_cor_ngd_smh_scs_crcxlh, rsfmri_cor_ngd_smh_scs_crcxrh, rsfmri_cor_ngd_smh_scs_hplh, rsfmri_cor_ngd_smh_scs_hprh, rsfmri_cor_ngd_smh_scs_pllh, rsfmri_cor_ngd_smh_scs_plrh, rsfmri_cor_ngd_smh_scs_ptlh, rsfmri_cor_ngd_smh_scs_ptrh, rsfmri_cor_ngd_smh_scs_thplh, rsfmri_cor_ngd_smh_scs_thprh, rsfmri_cor_ngd_smh_scs_vtdclh, rsfmri_cor_ngd_smh_scs_vtdcrh, rsfmri_cor_ngd_smm_scs_aalh, rsfmri_cor_ngd_smm_scs_aarh, rsfmri_cor_ngd_smm_scs_aglh, rsfmri_cor_ngd_smm_scs_agrh, rsfmri_cor_ngd_smm_scs_bs, rsfmri_cor_ngd_smm_scs_cdelh, rsfmri_cor_ngd_smm_scs_cderh, rsfmri_cor_ngd_smm_scs_crcxlh, rsfmri_cor_ngd_smm_scs_crcxrh, rsfmri_cor_ngd_smm_scs_hplh, rsfmri_cor_ngd_smm_scs_hprh, rsfmri_cor_ngd_smm_scs_pllh, rsfmri_cor_ngd_smm_scs_plrh, rsfmri_cor_ngd_smm_scs_ptlh, rsfmri_cor_ngd_smm_scs_ptrh, rsfmri_cor_ngd_smm_scs_thplh, rsfmri_cor_ngd_smm_scs_thprh, rsfmri_cor_ngd_smm_scs_vtdclh, rsfmri_cor_ngd_smm_scs_vtdcrh, rsfmri_cor_ngd_vta_scs_aalh, rsfmri_cor_ngd_vta_scs_aarh, rsfmri_cor_ngd_vta_scs_aglh, rsfmri_cor_ngd_vta_scs_agrh, rsfmri_cor_ngd_vta_scs_bs, rsfmri_cor_ngd_vta_scs_cdelh, rsfmri_cor_ngd_vta_scs_cderh, rsfmri_cor_ngd_vta_scs_crcxlh, rsfmri_cor_ngd_vta_scs_crcxrh, rsfmri_cor_ngd_vta_scs_hplh, rsfmri_cor_ngd_vta_scs_hprh, rsfmri_cor_ngd_vta_scs_pllh, rsfmri_cor_ngd_vta_scs_plrh, rsfmri_cor_ngd_vta_scs_ptlh, rsfmri_cor_ngd_vta_scs_ptrh, rsfmri_cor_ngd_vta_scs_thplh, rsfmri_cor_ngd_vta_scs_thprh, rsfmri_cor_ngd_vta_scs_vtdclh, rsfmri_cor_ngd_vta_scs_vtdcrh, rsfmri_cor_ngd_vs_scs_aalh, rsfmri_cor_ngd_vs_scs_aarh, rsfmri_cor_ngd_vs_scs_aglh, rsfmri_cor_ngd_vs_scs_agrh, rsfmri_cor_ngd_vs_scs_bs, rsfmri_cor_ngd_vs_scs_cdelh, rsfmri_cor_ngd_vs_scs_cderh, rsfmri_cor_ngd_vs_scs_crcxlh, rsfmri_cor_ngd_vs_scs_crcxrh, rsfmri_cor_ngd_vs_scs_hplh, rsfmri_cor_ngd_vs_scs_hprh, rsfmri_cor_ngd_vs_scs_pllh, rsfmri_cor_ngd_vs_scs_plrh, rsfmri_cor_ngd_vs_scs_ptlh, rsfmri_cor_ngd_vs_scs_ptrh, rsfmri_cor_ngd_vs_scs_thplh, rsfmri_cor_ngd_vs_scs_thprh, rsfmri_cor_ngd_vs_scs_vtdclh, rsfmri_cor_ngd_vs_scs_vtdcrh

**SI Table 5**. Factor loadings for the ten early-life adversity dimensions. Adapted from (Brieant et al., 2023), with permission from the author.

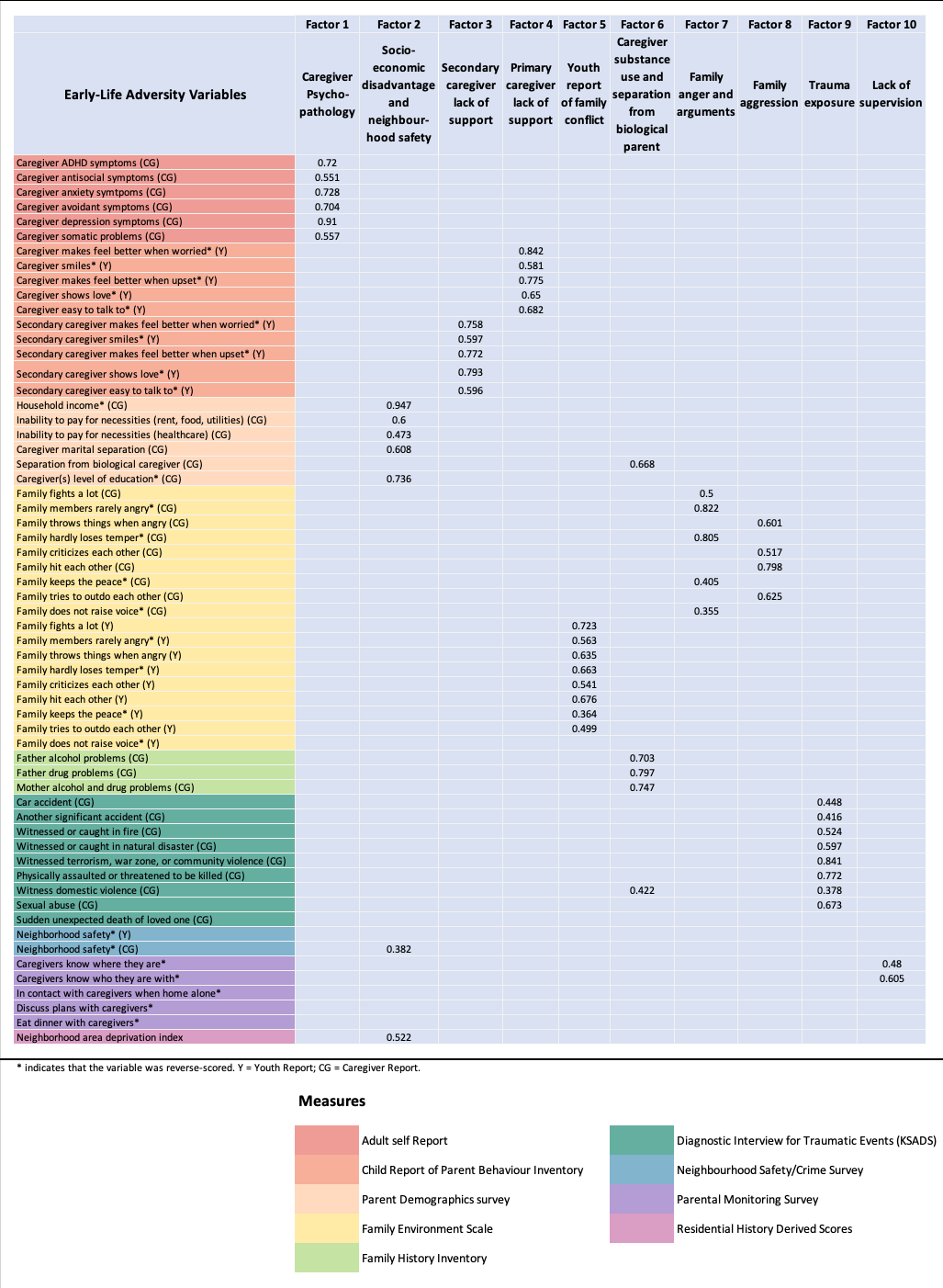

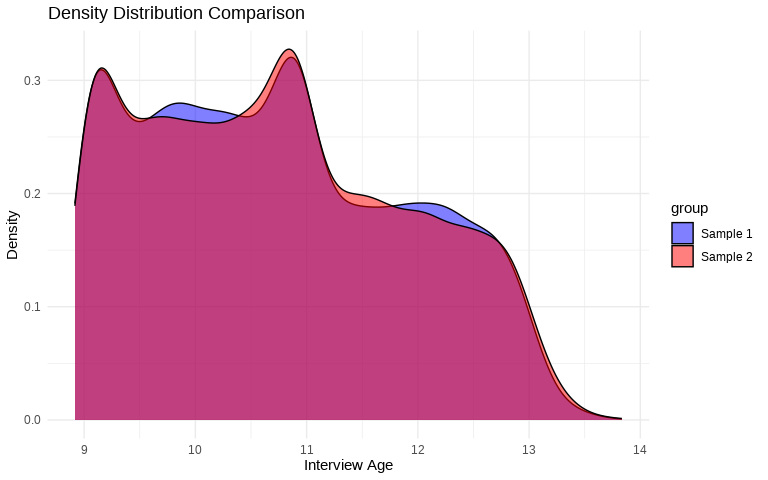

**SI Figure 4.** Age density distribution for T1-weighted-based training (Sample 1) and test (Sample 2) datasets.

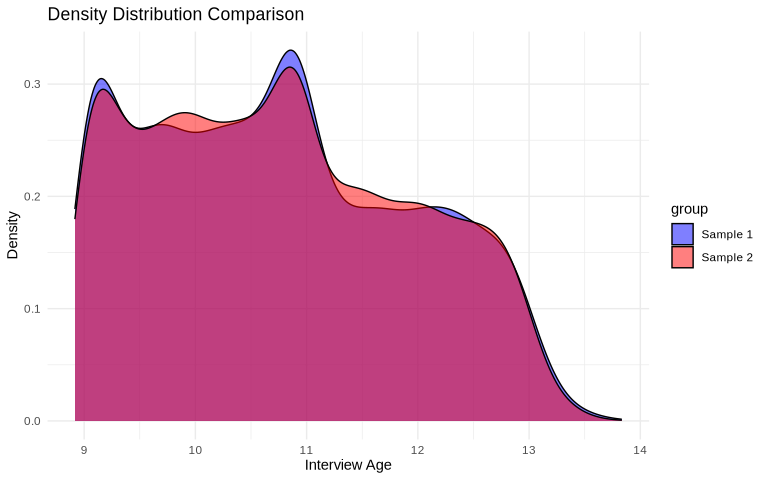

**SI Figure 5.** Age density distribution for DTI-based training (Sample 1) and test (Sample 2) datasets.

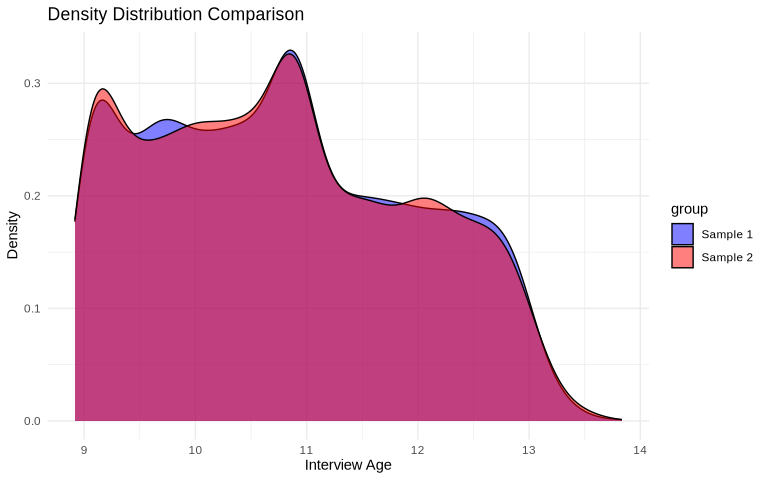

**SI Figure 6.** Age density distribution for rs-fMRI-based training (Sample 1) and test (Sample 2) datasets.

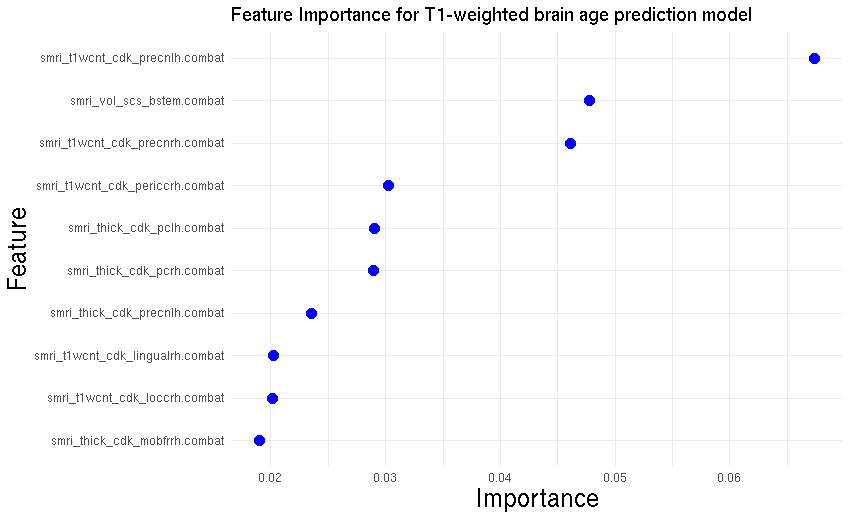

**SI Figure 7.** Showing top ten T1-weighted brain age model feature importance scores. Range of 0.01-0.07 represents a relative contribution of 1-7% importance. Top three features include smri_t1wcnt_cdk_precnlh (and rh), representing grey-white contrast of T1 weighted image for APARC ROI precentral; and smri_vol_scs_bstem, representing volume in mm^3 of ASEG ROI brain-stem.

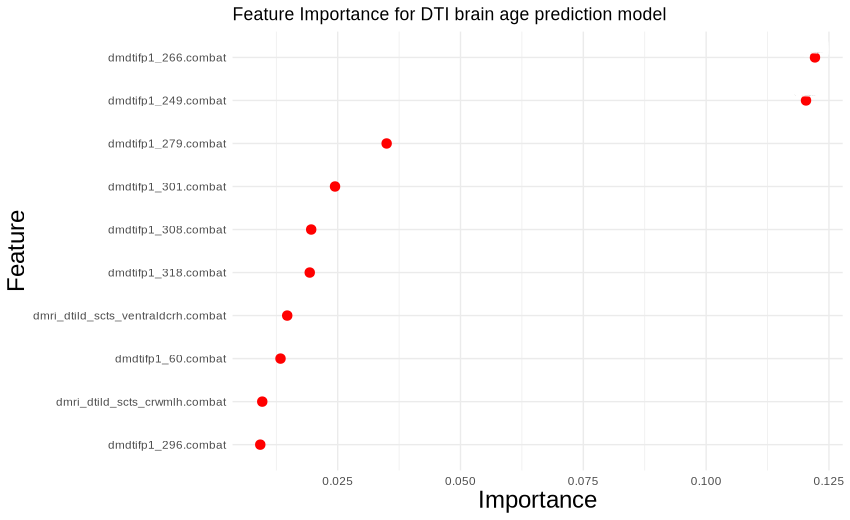

**SI Figure 8.** Showing top ten DTI brain age model feature importance scores. Range of 0.01-0.3 represents a relative contribution of 1-13% importance. The top three features include dmdtifp1_266, dmdtifp1_249, dmdtifp1_279, which represent the mean diffusivity within ASEG ROI right-pallidum, the mean diffusivity within ASEG ROI left-pallidum, and the average longitudinal diffusion coefficient within ASEG ROI left-pallidum, respectively.

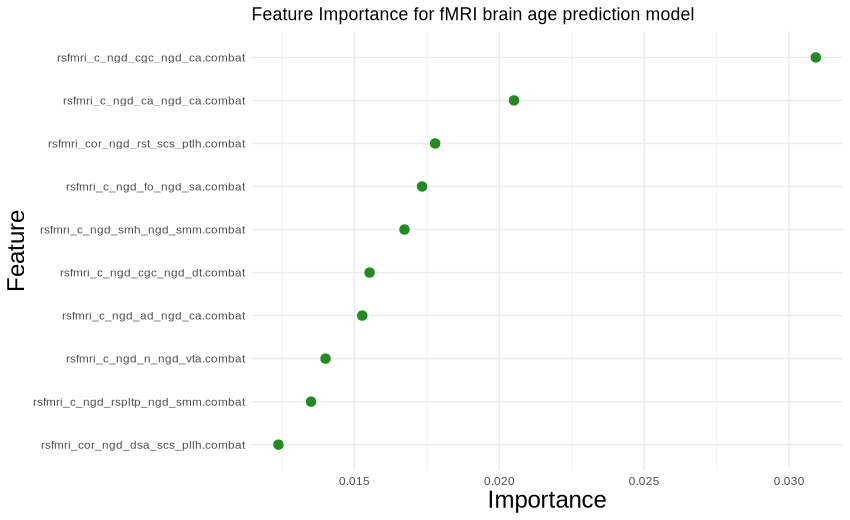

**SI Figure 9.** Showing top ten fMRI brain age model feature importance scores. Range of 0.01-0.04 represents a relative contribution of 1-4% importance. The top three features include rsfmri_c_ngd_cgc_ngd_ca, the average correlation between cingulo-opercular network and cingulo-parietal network; rsfmri_c_ngd_ca_ngd_ca, the average correlation between cingulo-parietal network and cingulo-parietal network; and rsfmri_cor_ngd_rst_scs_ptlh, the average correlation between retrosplenial temporal network and ASEG ROI left-putamen.

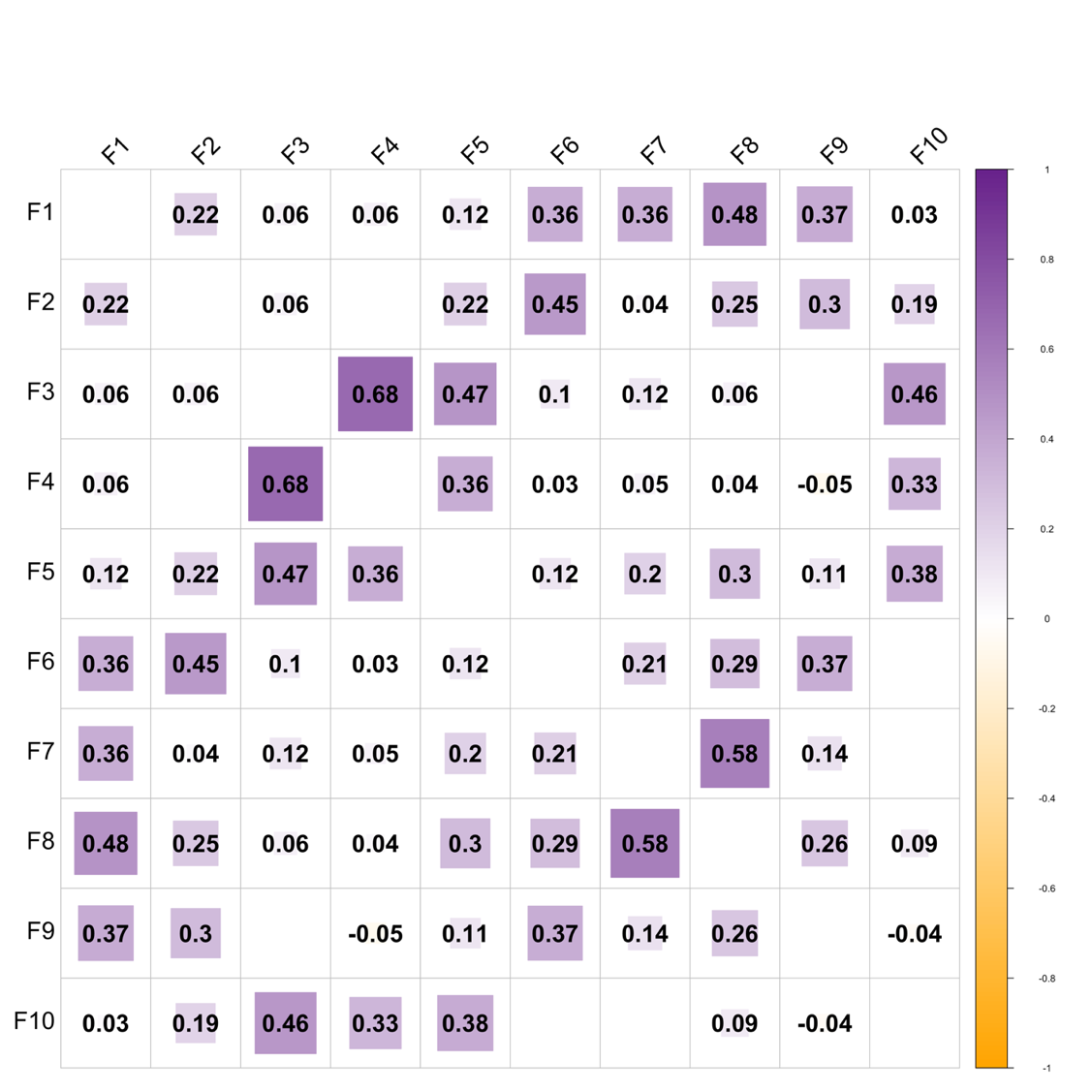

**SI Figure 10.** Correlation matrix showing the associations between each of the 10 dimensions of ELA.

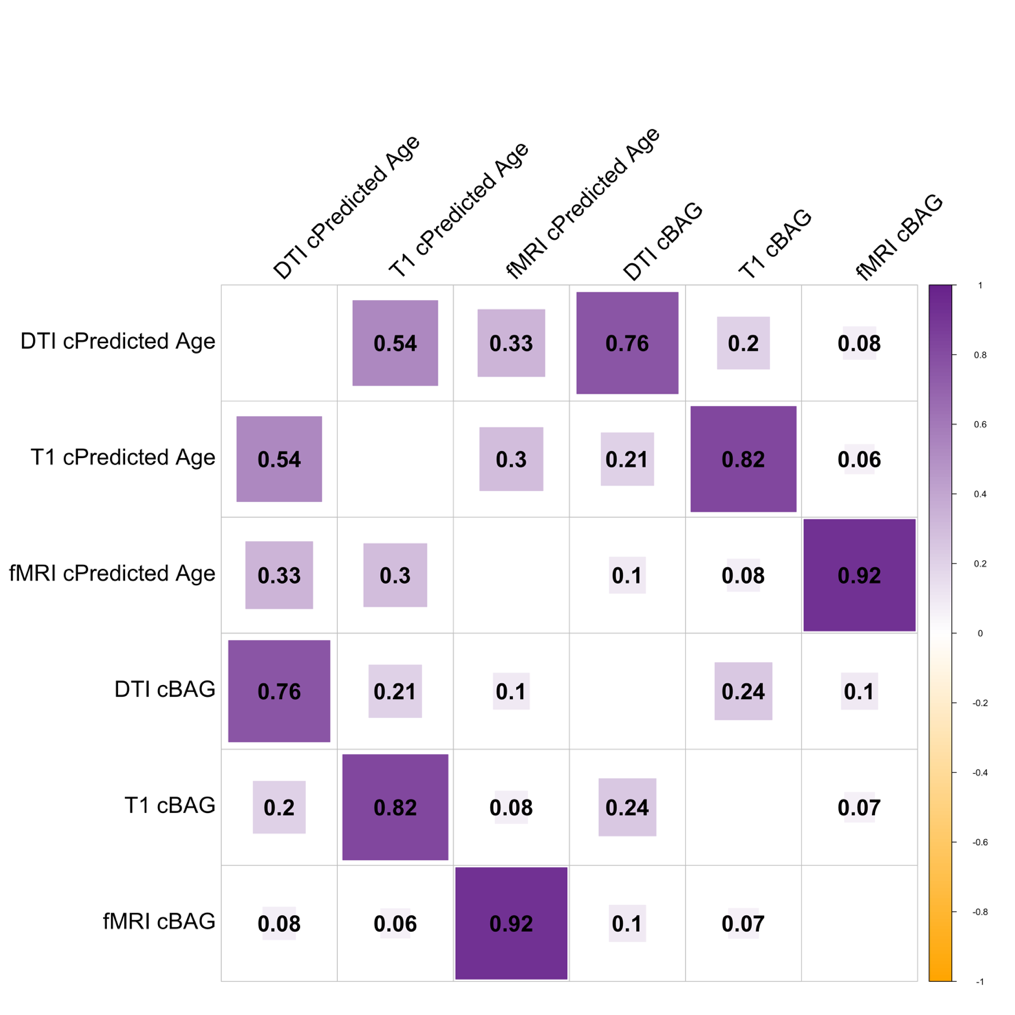

**SI Figure 11.** Correlation matrix showing the associations between modality-specific residualised brain predicted age and residualised brain age gap (cBAG).

| **SI Table 6.** **Bayes Factor (BF).** Showing evidence ratio interpretations. Values of 1 can be interpreted as no evidence in either direction, with the following values indicating weight of evidence towards the alternative hypothesis: 0.3-1 (anecdotal), 0.1-0.3 (moderate), 0.03-0.1 (strong), 0.01- 0.03 (very strong), <0.01 (extreme). Contrarily, the following values indicate weight of evidence towards the null hypothesis: 1-3 (anecdotal), 3-10 (moderate), 10-30 (strong), 30-100 (very strong), >100 (extreme). | | | |
| --- | --- | --- | --- |
| Bayes factor BF_12_ | | | Interpretation |
|  | > | 100 | Extreme evidence for M_1_ |
| 30 | - | 100 | Very strong evidence for M_1_ |
| 10 | - | 30 | Strong evidence for M_1_ |
| 3 | - | 10 | Moderate evidence for M_1_ |
| 1 | - | 3 | Anecdotal evidence for M_1_ |
|  | 1 |  | No evidence |
| 1/3 | - | 1 | Anecdotal evidence for M_2_ |
| 1/10 | - | 1/3 | Moderate evidence for M_2_ |
| 1/30 | - | 1/10 | Strong evidence for M_2_ |
| 1/100 | - | 1/30 | Very strong evidence for M_2_ |
|  | < | 1/100 | Extreme evidence for M_2_ |

| **SI Table 7**. Bayesian multilevel model results for T1-weighted associations with ELA dimensions. Table shows results of main effects and interaction effects between T1 BAG and ELA dimensions and T1 BAG and ELA dimensions x Timepoint, respectively. Evidence ratios under 1 indicate evidence for the alternative hypothesis and values over 1 indicate evidence for the null hypothesis. Estimates indicate directionality of evidence (positive/negative association). See SI Table 2 for precise interpretation of evidence ratio values. | | | | | |
| --- | --- | --- | --- | --- | --- |
| Modality | ELA | Estimate | Lower95 | Upper95 | Evidence  ratio |
| T1 cBAG | F1 | -0.03 | -0.08 | 0.02 | 2.16 |
| T1 cBAG | F2 | 0.13 | 0.08 | 0.19 | 0 |
| T1 cBAG | F3 | -0.05 | -0.12 | 0.01 | 0.91 |
| T1 cBAG | F4 | -0.08 | -0.14 | -0.02 | 0.17 |
| T1 cBAG | F5 | 0.01 | -0.05 | 0.08 | 2.69 |
| T1 cBAG | F6 | 0.05 | -0.02 | 0.12 | 1.14 |
| T1 cBAG | F7 | -0.01 | -0.08 | 0.05 | 2.85 |
| T1 cBAG | F8 | 0 | -0.07 | 0.07 | 2.88 |
| T1 cBAG | F9 | 0.05 | -0.03 | 0.13 | 1.07 |
| T1 cBAG | F10 | -0.02 | -0.08 | 0.05 | 2.65 |
| T1 cBAG | TP:F1 | 0.03 | -0.02 | 0.09 | 1.67 |
| T1 cBAG | TP:F2 | 0.01 | -0.05 | 0.06 | 3.27 |
| T1 cBAG | TP:F3 | -0.04 | -0.11 | 0.02 | 1.21 |
| T1 cBAG | TP:F4 | -0.03 | -0.09 | 0.03 | 2.21 |
| T1 cBAG | TP:F5 | -0.04 | -0.1 | 0.02 | 1.45 |
| T1 cBAG | TP:F6 | 0.02 | -0.05 | 0.09 | 2.27 |
| T1 cBAG | TP:F7 | -0.01 | -0.07 | 0.06 | 2.96 |
| T1 cBAG | TP:F8 | 0.03 | -0.03 | 0.09 | 2.02 |
| T1 cBAG | TP:F9 | 0 | -0.07 | 0.08 | 2.72 |
| T1 cBAG | TP:F10 | -0.09 | -0.15 | -0.03 | 0.08 |
| ***Note:*** *cBAG represents residualised (age-bias corrected) brain age gap scores.* | | | | | |

| **SI Table 8.** Bayesian multilevel model results for DTI associations with ELA dimensions. Table shows results of main effects and interaction effects between DTI BAG and ELA dimensions and DTI BAG and ELA dimensions x Timepoint, respectively. Evidence ratios under 1 indicate evidence for the alternative hypothesis and values over 1 indicate evidence for the null hypothesis. Estimates indicate directionality of evidence (positive/negative association). See SI Table 2 for precise interpretation of evidence ratio values. | | | | | |
| --- | --- | --- | --- | --- | --- |
| Modality | ELA | Estimate | Lower95 | Upper95 | Evidence  ratio |
| DTI cBAG | F1 | 0.01 | -0.03 | 0.05 | 4.09 |
| DTI cBAG | F2 | 0.06 | 0.01 | 0.1 | 0.27 |
| DTI cBAG | F3 | 0.02 | -0.04 | 0.07 | 3.22 |
| DTI cBAG | F4 | 0.02 | -0.04 | 0.07 | 3.15 |
| DTI cBAG | F5 | 0.01 | -0.04 | 0.06 | 3.56 |
| DTI cBAG | F6 | 0.04 | -0.02 | 0.1 | 1.32 |
| DTI cBAG | F7 | -0.02 | -0.07 | 0.03 | 3.05 |
| DTI cBAG | F8 | -0.03 | -0.09 | 0.02 | 1.9 |
| DTI cBAG | F9 | 0.04 | -0.02 | 0.1 | 1.54 |
| DTI cBAG | F10 | 0 | -0.06 | 0.05 | 3.63 |
| DTI cBAG | TP:F1 | 0.07 | 0.02 | 0.13 | 0.09 |
| DTI cBAG | TP:F2 | 0.02 | -0.04 | 0.07 | 2.86 |
| DTI cBAG | TP:F3 | 0.01 | -0.05 | 0.07 | 3.14 |
| DTI cBAG | TP:F4 | 0.01 | -0.05 | 0.08 | 2.91 |
| DTI cBAG | TP:F5 | 0 | -0.06 | 0.06 | 3.16 |
| DTI cBAG | TP:F6 | 0.05 | -0.02 | 0.12 | 1.2 |
| DTI cBAG | TP:F7 | 0.04 | -0.02 | 0.1 | 1.44 |
| DTI cBAG | TP:F8 | 0.03 | -0.03 | 0.1 | 1.78 |
| DTI cBAG | TP:F9 | 0.03 | -0.04 | 0.1 | 1.93 |
| DTI cBAG | TP:F10 | -0.02 | -0.08 | 0.05 | 2.58 |

| **SI Table 9.** Bayesian multilevel model results for rs-fMRI associations with ELA dimensions. Table shows results of main effects and interaction effects between rs-fMRI BAG and ELA dimensions and rs-fMRI BAG and ELA dimensions x Timepoint, respectively. Evidence ratios under 1 indicate evidence for the alternative hypothesis and values over 1 indicate evidence for the null hypothesis. Estimates indicate directionality of evidence (positive/negative association). See SI Table 2 for precise interpretation of evidence ratio values. | | | | | |
| --- | --- | --- | --- | --- | --- |
| Modality | ELA | Estimate | Lower95 | Upper95 | Evidence  ratio |
| rs-fMRI cBAG | F1 | 0.07 | -0.02 | 0.14 | 0.63 |
| rs-fMRI cBAG | F2 | 0.18 | 0.1 | 0.27 | 0 |
| rs-fMRI cBAG | F3 | -0.05 | -0.15 | 0.04 | 1.19 |
| rs-fMRI cBAG | F4 | -0.1 | -0.19 | -0.01 | 0.27 |
| rs-fMRI cBAG | F5 | -0.05 | -0.15 | 0.04 | 1.19 |
| rs-fMRI cBAG | F6 | 0.15 | 0.05 | 0.26 | 0.03 |
| rs-fMRI cBAG | F7 | 0.03 | -0.06 | 0.13 | 1.57 |
| rs-fMRI cBAG | F8 | 0.08 | -0.02 | 0.18 | 0.66 |
| rs-fMRI cBAG | F9 | 0.15 | 0.03 | 0.25 | 0.07 |
| rs-fMRI cBAG | F10 | 0 | -0.1 | 0.1 | 1.99 |
| rs-fMRI cBAG | TP:F1 | 0.01 | -0.08 | 0.1 | 2.07 |
| rs-fMRI cBAG | TP:F2 | 0.03 | -0.07 | 0.13 | 1.71 |
| rs-fMRI cBAG | TP:F3 | -0.08 | -0.18 | 0.02 | 0.62 |
| rs-fMRI cBAG | TP:F4 | -0.06 | -0.15 | 0.05 | 1.05 |
| rs-fMRI cBAG | TP:F5 | -0.08 | -0.18 | 0.02 | 0.58 |
| rs-fMRI cBAG | TP:F6 | 0.09 | -0.02 | 0.19 | 0.51 |
| rs-fMRI cBAG | TP:F7 | 0 | -0.1 | 0.1 | 2 |
| rs-fMRI cBAG | TP:F8 | 0.03 | -0.07 | 0.13 | 1.64 |
| rs-fMRI cBAG | TP:F9 | 0.01 | -0.1 | 0.12 | 1.74 |
| rs-fMRI cBAG | TP:F10 | -0.02 | -0.12 | 0.09 | 1.73 |

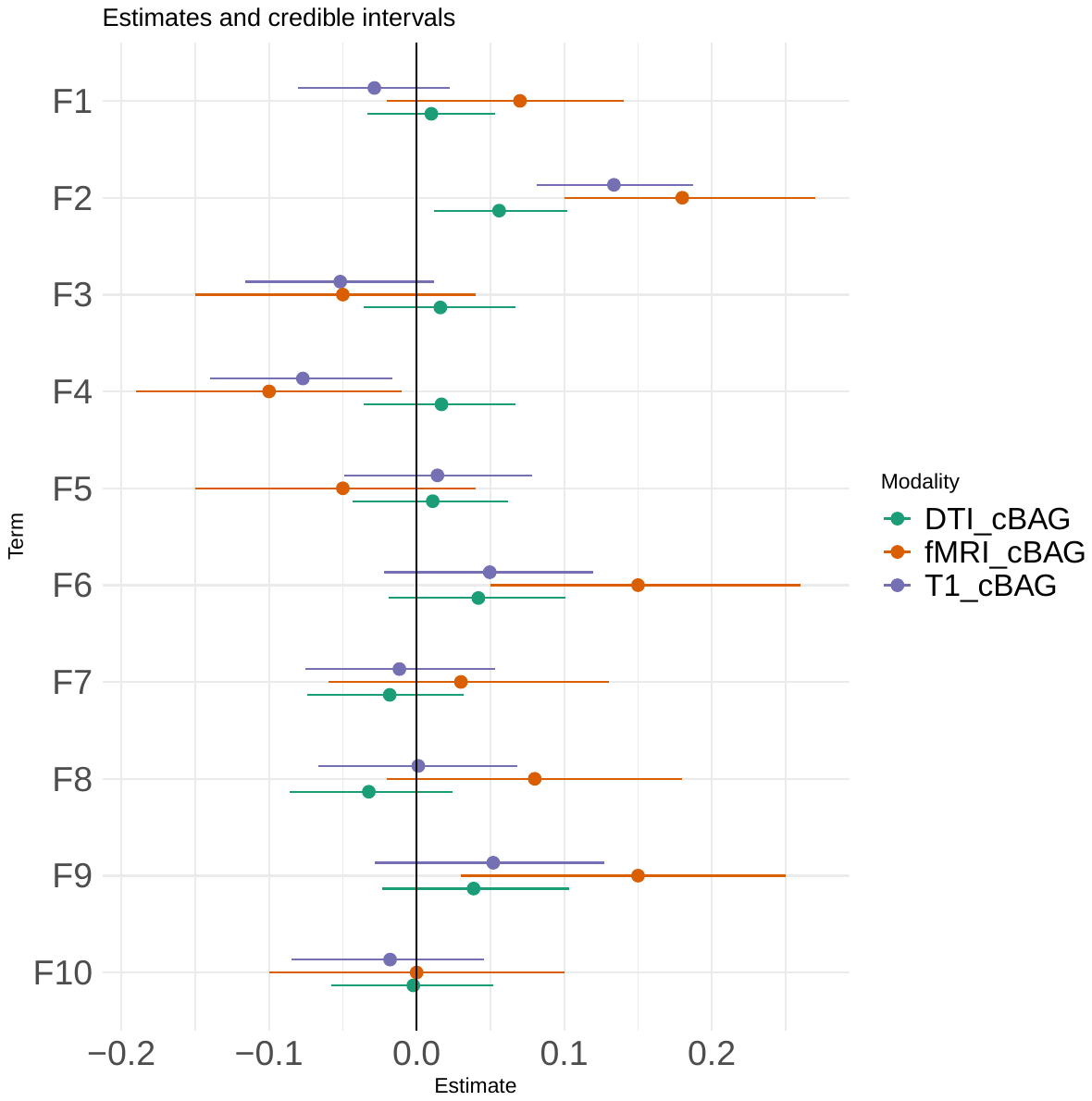

**SI Figure 12.** **Estimate credible intervals.** Showing the association (main effect) between modality-specific BAG and ELA Dimensions. Figure shows estimates with 95% credible intervals. T1 associations are represented by purple, DTI associations represented by green, and rs-fMRI represented by orange.

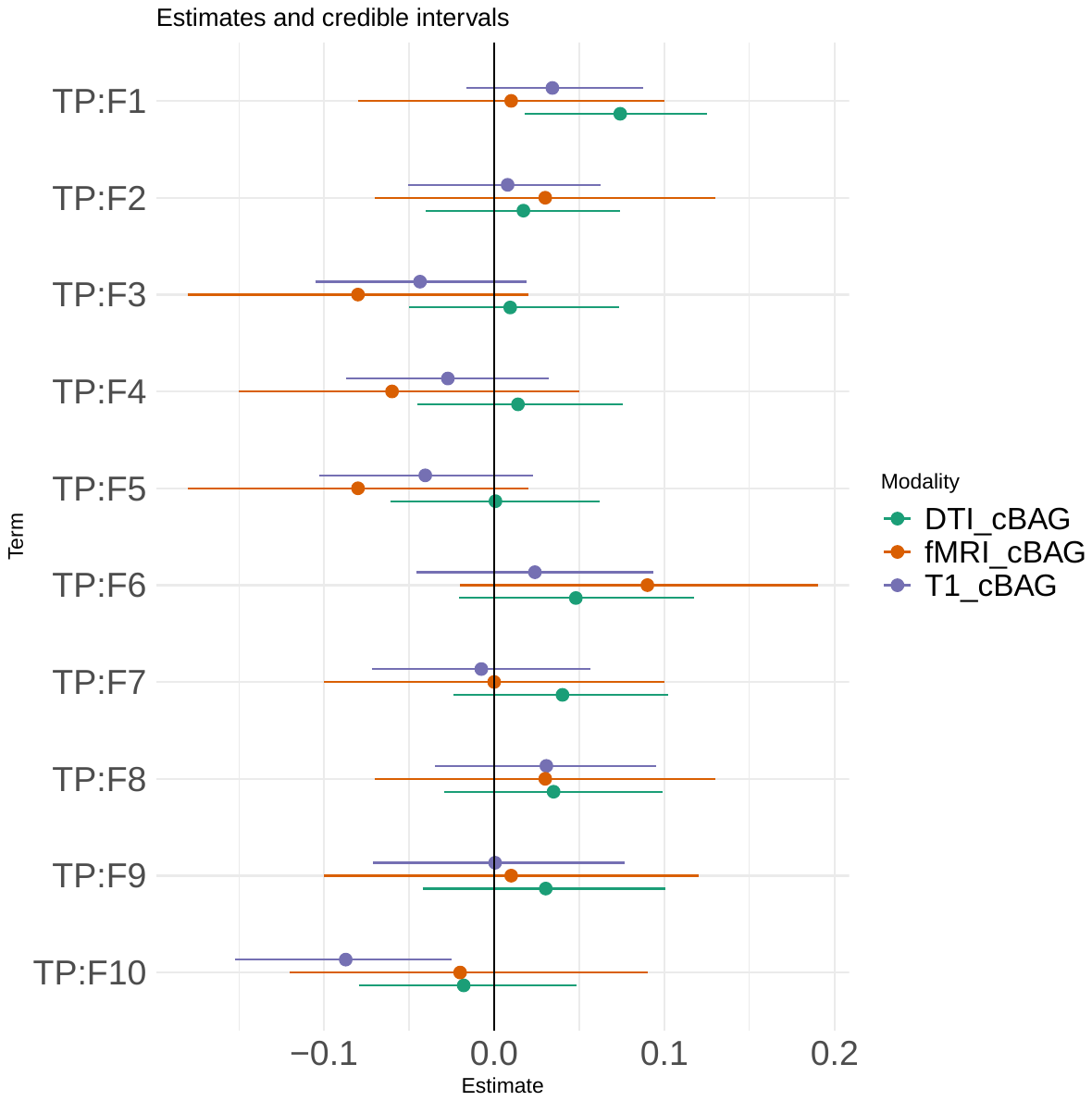

**SI Figure 13.** **Estimate credible intervals**. Showing the association (interaction effect) between modality-specific BAG and ELA*Timepoint. Figure shows estimates with 95% credible intervals. T1 associations are represented by purple, DTI associations represented by green, and rs-fMRI represented by orange.
